# Senescent cells induce vascular MHC II to recruit CD4^+^ T cells and drive inflammation in aging adipose tissue

**DOI:** 10.64898/2026.05.13.724992

**Authors:** Qijing Xie, Tzuhua D. Lin, SoRi Jang, Calvin Jan, Amirali Selahi, Kendra Nyberg, Agnieszka A. Wendorff, Kayley Hake, Austin E.Y.T. Lefebvre, Meng Jin, Andrea Di Francesco, Maria Pokrovskii, Helen Tauc-Adrian, Cynthia Kenyon

## Abstract

Adipose tissue exhibits pronounced inflammation during aging, yet the mechanisms sustaining this chronic state are not well understood. By creating an atlas integrating histology, single-nucleus transcriptomics and flow cytometry across the murine lifespan, we find that age-associated inflammation is distinct from the obesity-like inflammatory profile observed at mid-life. Specifically, age-associated inflammation is characterized by a potent interferon-gamma (IFNγ) response signature and the accumulation of T cells. We demonstrate that senescent cells act as an upstream trigger, indirectly initiating an IFNγ response that upregulates vascular MHC II to promote extravasation of CD4^+^ T cells into the aging tissue. These recruited T cells then act as a critical source of IFNγ, thereby perpetuating a positive feedback loop that maintains chronic immune infiltration and tissue inflammation. These results provide a multi-modal view of adipose aging and identify a mechanism that sustains its age-associated inflammation.

## Main

Adipose tissue is increasingly recognized as a critical regulator of the aging process^1–3^. As a major energy reservoir and an important immunoendocrine organ, it maintains energy homeostasis and modulates systemic inflammatory tone throughout the lifespan^4–7^. Disruptions in adipose tissue homeostasis––whether through excess accumulation or significant loss––are associated with an increased risk of all-cause mortality and a spectrum of age-related diseases^8–12^. Conversely, interventions that preserve or restore adipose tissue homeostasis often exert a beneficial impact on healthspan and lifespan^13–17^. Understanding how adipose tissue ages may therefore provide critical insights into maintaining health during aging.

On the macroscopic level, adipose tissue undergoes significant remodeling during the natural aging process. In humans and *ad libitum*-fed mice, adipose tissue mass usually follows a biphasic trajectory, increasing from young to middle age before undergoing a progressive decline toward the end of life^18–20^. In addition, lipids have been reported to redistribute from subcutaneous to visceral and ectopic depots with age––a shift associated with exacerbated metabolic dysfunction and increased disease risk^1,2^.

At the molecular and cellular levels, adipose tissue develops pronounced inflammation with age^21,22^. This inflammatory state is frequently attributed to the stress induced by tissue expansion, which triggers macrophage infiltration and the elevated production of pro-inflammatory cytokines^23,24^. However, intrinsic aging processes, such as the accumulation of senescent cells, are also known to promote tissue inflammation^25^. Because adipose mass follows a biphasic trajectory, the use of only two discrete timepoints (“young” versus “aged”) in most studies may not provide the temporal resolution required to follow this biphasic dynamic and thereby decouple mass-associated from age-associated signals. Consequently, it remains unknown whether age-related inflammation is simply a consequence of unresolved inflammation from the mid-life adiposity peak, or if a distinct, age-specific mechanism is responsible for this chronic state.

In this study, we integrated histology, single-nucleus RNA sequencing, and flow cytometry to map the structural, molecular, and cellular landscape of adipose tissue across five timepoints spanning the murine lifespan. Our analysis revealed two distinct waves of inflammation: an initial wave that mirrored the inflammation associated with adipose tissue expansion and a second, age-associated wave characterized by a potent interferon-gamma (IFNγ) response signature. In exploring the mechanisms underlying this age-associated wave, we utilized *in vivo* senescent cell transplantation and *in vitro* modeling to show that the accumulation of senescent cells led to the upregulation of MHC II on the vasculature, which facilitated a non-canonical, antigen-dependent extravasation of CD4^+^ T cells into the aging tissue. We further demonstrated that these CD4^+^ T cells were required for the IFNγ response signature and for the maintenance of age-related immune cell infiltrates. Together, these findings established a multi-modal atlas of adipose tissue aging and identified a specific cellular circuit that maintains age-related inflammation.

## Results

### Age-related adipose tissue remodeling

To better describe the adipose aging trajectory, we analyzed subcutaneous inguinal (SAT) and visceral perigonadal (VAT) white adipose tissue from *ad libitum*-fed C57BL/6J mice across five ages (Fig. 1a). Adipose tissue weight followed a biphasic trajectory in both sexes, albeit with distinct patterns: males exhibited a peak at 18 months before declining, whereas females reached a plateau that was maintained from 12 through 24 months (Fig. 1b and Extended Data Fig. 1a,b). Because the pronounced 18-month inflection point in males allows for a clearer distinction between the impacts of tissue expansion and chronological aging, we focused our subsequent investigation on male mice (Supplementary Table 1).

**Fig. 1.**
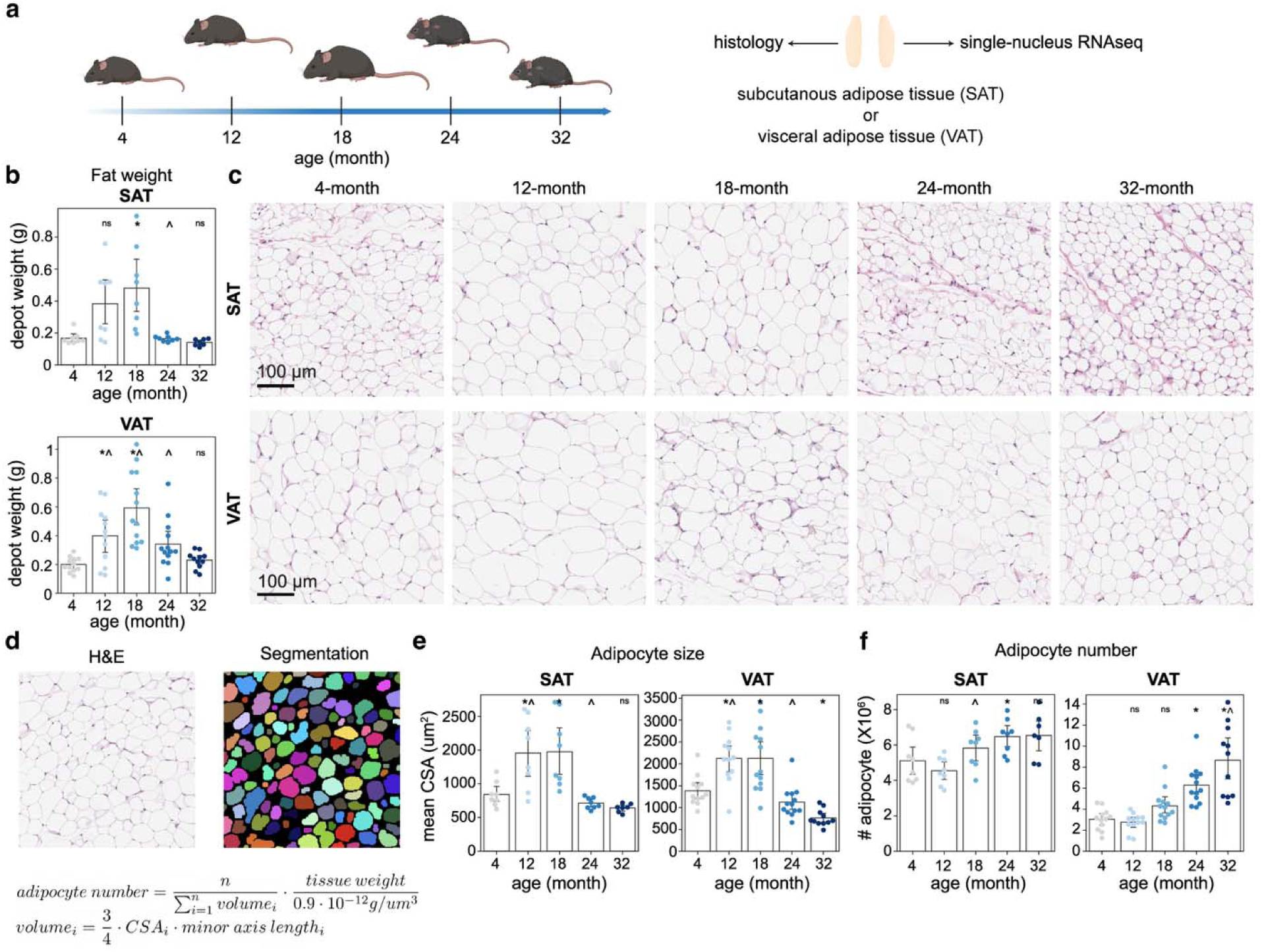
| Depot-specific and age-associated structural remodeling of adipose tissue. **a**, Schematic overview of the experimental design for characterizing subcutaneous (SAT) and visceral (VAT) adipose tissue across the murine lifespan. **b**, Weight of SAT (*n* = 7–8 per group) and VAT (*n* = 12–13 per group) across five age groups. **c**, Representative images of Hematoxylin and Eosin (H&E)-stained SAT and VAT sections across five age groups. **d**, Image analysis strategy for the quantification of mean cross-sectional area (CSA) and estimation of total number of adipocytes from histological sections. **e**, **f**, Mean CSA (**e**) and estimated number of adipocytes (**f**) in SAT (*n* = 7–8 per group) and VAT (*n* = 11–13 per group) across five age groups. In each bar plot (**b, e** and **f**), data are presented as mean ± 95% confidence intervals, with each dot representing an individual mouse. Statistical significance was determined by one-way ANOVA with Tukey’s post hoc test or Welch’s two-sided t-test when variances were not equal. Asterisks (*) indicate p-value < 0.05 compared to 4-month-old mice, and carets (^) indicate p-value < 0.05 compared to the preceding age group.

We quantified histology images to determine how adipocyte size and number contributed to changes in adipose tissue weight with age (Fig. 1c,d). We found that adipose tissue undergoes two distinct phases of expansion, comprising an initial period of adipocyte hypertrophy (cell size increase) from 4 to 12 months followed by a transition to hyperplastic growth (cell number increase) between 12 and 18 months (Fig. 1 e,f and Extended Data Fig. 1 c,d). This hypertrophy-to-hyperplasia transition was reminiscent of the cellular events observed during the progression of obesity^26,27^. Coinciding with this expansion phase, we observed the accumulation of lipid vacuoles in the liver (Extended Data Fig. 1e,f). Notably, the area of these liver lipid vacuoles correlated with adipose tissue weight but not with age (Extended Data Fig. 1g). These data suggested that male mice developed a mild, obesity-like phenotype during mid-life.

Following the peak at 18 months, weights of the two depots declined at different rates that correlated with distinct cellular profiles (Fig. 1b). SAT weight dropped sharply between 18 and 24 months as adipocyte size decreased (Fig. 1b,c,e). SAT adipocyte size and tissue weight then remained stable through the end-of-life. In contrast, VAT adipocytes underwent continued atrophy and decreased to approximately half their 4-month-old size at 32 months (Fig. 1b,c,e). Despite this pronounced decrease in cell size, VAT exhibited sustained hyperplasia through late life (Fig. 1f). This continuous production of new adipocytes in the VAT potentially offset the reduction in individual cell size, resulting in a slower decline in tissue weight.

In summary, these findings revealed that mid-life adipose tissue expansion recapitulated an obesity-like state, whereas the subsequent late-life decline in mass was characterized by depot-specific cellular remodeling.

### An integrated transcriptomic atlas of aging adipose tissue

To identify molecular programs underlying these structural changes and explore other age-associated biological shifts, we performed single-nucleus RNA sequencing on SAT and VAT collected from the mice described above (Fig. 1a). Although other single-cell transcriptional atlases for aging mouse adipose tissue exist^28–30^, our multi-modal approach allowed us to match gene expression with the structural changes we observed. Using a split-pool combinatorial barcoding approach, we obtained transcriptomes characteristic of white adipose tissue^31–33^, including adipocytes, adipose stem and progenitor cells (ASPCs), endothelial cells and immune cells (Fig. 2a and Extended Data Fig. 1h–j).

**Fig. 2.**
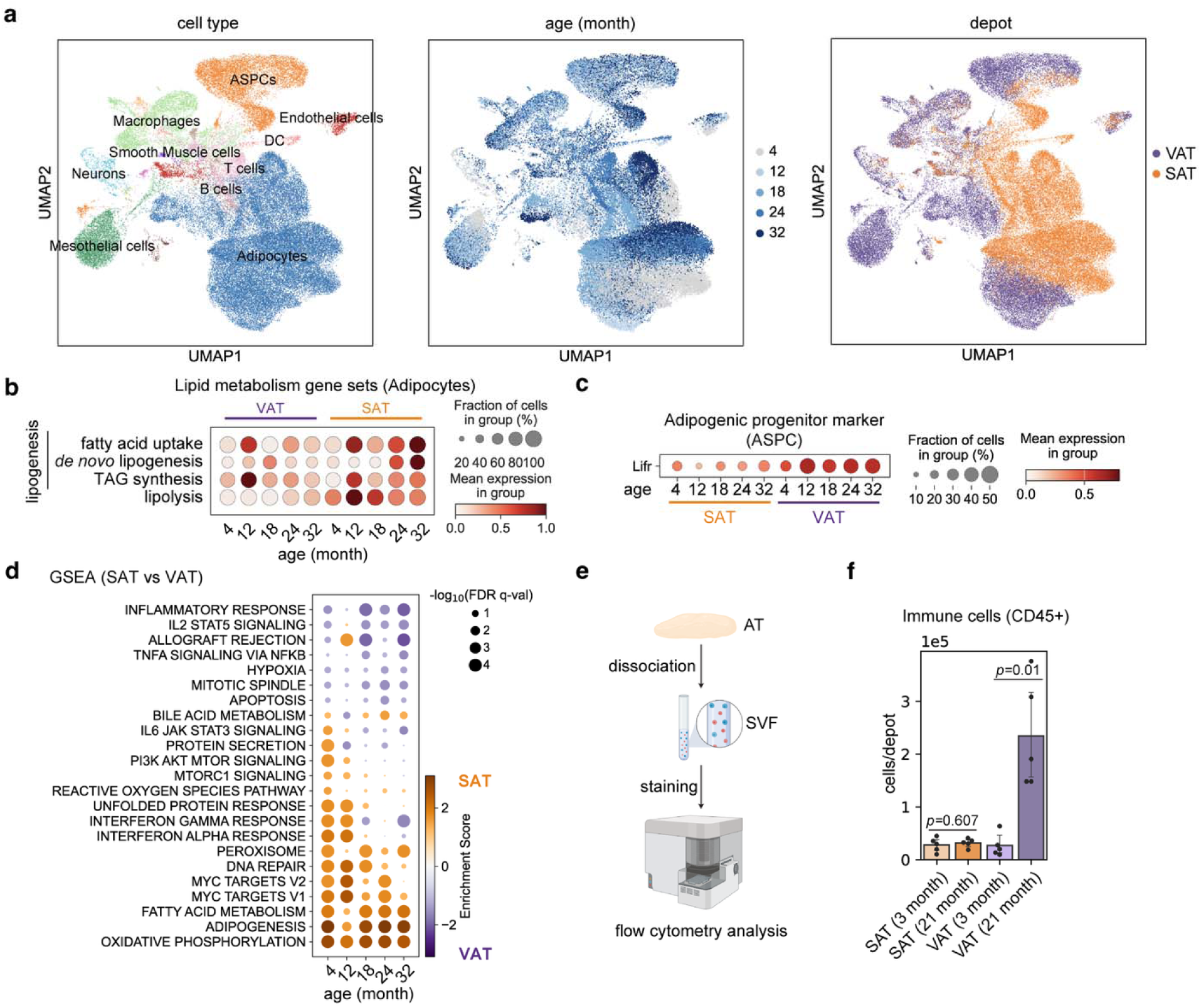
|. Aging adipose tissue atlas reveals depot-specific transcriptomic and immunological signatures. **a**, Uniform Manifold Approximation and Projection (UMAP) of sequenced nuclei from SAT and VAT, colored by major cell types (left), age (middle) and depot (right). **b**, Expression of lipid metabolism gene sets in SAT and VAT adipocytes at different ages. Expression of individual genes is detailed in Extended Data Fig. 2b. **c**, Dot plot showing the expression of *Lifr* in ASPCs. **d**, Gene set enrichment analysis (GSEA) revealed transcriptional differences between SAT and VAT at each age. A positive enrichment score (orange) indicates higher expression in SAT. Dot size corresponds to the false discovery rate (FDR) q-value. **e**, Experimental workflow for flow cytometry analysis of the stromal vascular fraction (SVF) of adipose tissue. **f**, Total immune cell number in the SVF from young (3-month-old) and old (21-month-old) SAT and VAT (*n* = 5 per group). Data are presented as mean ± 95% confidence intervals, with each dot representing an individual mouse. Statistical significance was determined by Welch’s two-sided t-test.

We next asked if changes in gene expression could suggest explanations for the depot-specific, age-related changes in adipocyte size and number. Indeed, the changes in adipocyte size matched the expression of genes involved in lipid storage and breakdown (Extended Data Fig. 2a)^34^. During the hypertrophic growth phase from 4 to 12 months, adipocytes in both depots upregulated lipogenesis genes, including those involved in fatty acid uptake and triacylglyceride (TAG) synthesis (Fig. 2b and Extended Data Fig. 2b). During the phase of adipocyte size decline from 18 to 32 months, the two depots diverged: VAT adipocytes maintained low expression of lipogenesis genes and upregulated lipolysis genes, whereas SAT adipocytes upregulated lipogenesis genes and downregulated lipolysis genes, matching the continued decrease of adipocyte size in VAT and the stabilized adipocyte size in SAT (Fig. 1e and 2b).

Similarly, the transcriptomic profile of ASPCs provided a potential explanation for the sustained increase in VAT adipocyte number through late life. We found that *Lifr*, a functional marker for the highly adipogenic progenitor population that emerges in aging VAT^35^, was upregulated in VAT ASPCs at 12 months and remained elevated through 32 months (Fig. 2c). This sustained expression aligned with the continuous production of new adipocytes identified in our histological analysis (Fig. 1f).

Together, this multi-modal atlas correlated specific transcriptomic changes with the structural remodeling we observed. Beyond these physical changes, the atlas also suggested that cells from SAT and VAT might follow different age-dependent transcriptional shifts (Fig. 2a), leading us to investigate the specific transcriptomic signatures of each depot.

### Depot-specific aging signatures

To identify depot-specific transcriptomic signatures, we compared the pseudobulk expression profiles of the two depots at each age using Gene Set Enrichment Analysis (GSEA) (Supplementary Table 2). We found that SAT was consistently enriched for “oxidative phosphorylation”, “adipogenesis” and “fatty acid metabolism” gene sets across all five ages (Fig. 2d). Supporting this result, the COMPASS algorithm^36^ predicted that SAT adipocytes had significantly higher lipid metabolism activity compared to VAT adipocytes, alongside various other metabolic pathways (Extended Data Fig. 2c,d).

In contrast, VAT displayed a progressively inflammatory profile. The VAT showed a subtle enrichment for “inflammatory response” and “TNF-α signaling via NF-κB” gene sets at 4 months, and this difference became more pronounced with age (Fig. 2d). Other key inflammatory pathways, such as the “interferon gamma (IFNγ) response” and “IL6 JAK STAT3 signaling” were initially more prominent in SAT at 4 months but became progressively enriched in VAT as mice aged. This inflammatory shift was pervasive across the visceral depot, as GSEA performed on age-associated genes within each cell type revealed a more pronounced upregulation of these pathways in VAT compared to their SAT counterparts (Extended Data Fig. 2e).

To investigate whether this inflammatory transcriptomic signature in VAT was accompanied by immune cell infiltration, we performed flow cytometry analysis on SAT and VAT from 4- and 21-month-old mice (Fig. 2e, Extended Data Fig. 2f, and Supplementary Table 3). We selected 21 months as a representative aged timepoint because the divergence in inflammatory gene expression between these depots was observed at all timepoints from 18 months onward (Fig. 2d). Consistent with our transcriptomic findings, the total number of immune cells increased significantly in VAT but remained unchanged in SAT (Fig. 2f). This increase in both inflammatory gene expression and immune cell number, two hallmarks of inflammation, led us to focus on VAT for our subsequent mechanistic analyses of age-associated inflammation.

### Two unique waves of inflammation

Since the obesity-like expansion of adipose tissue occurred only through middle age (Fig. 1), we reasoned that age-associated inflammation may be a result of either unresolved inflammation from this early peak in metabolic stress or the development of a distinct, age-associated mechanism. To investigate whether inflammation occurred with the peak of adipose tissue weight or if it was upregulated continuously with age, for each major cell type, we selected genes with significant expression changes across the five timepoints and clustered them into six distinct temporal trajectories (Fig. 3a and Extended Data Fig. 3a,b).

**Fig. 3.**
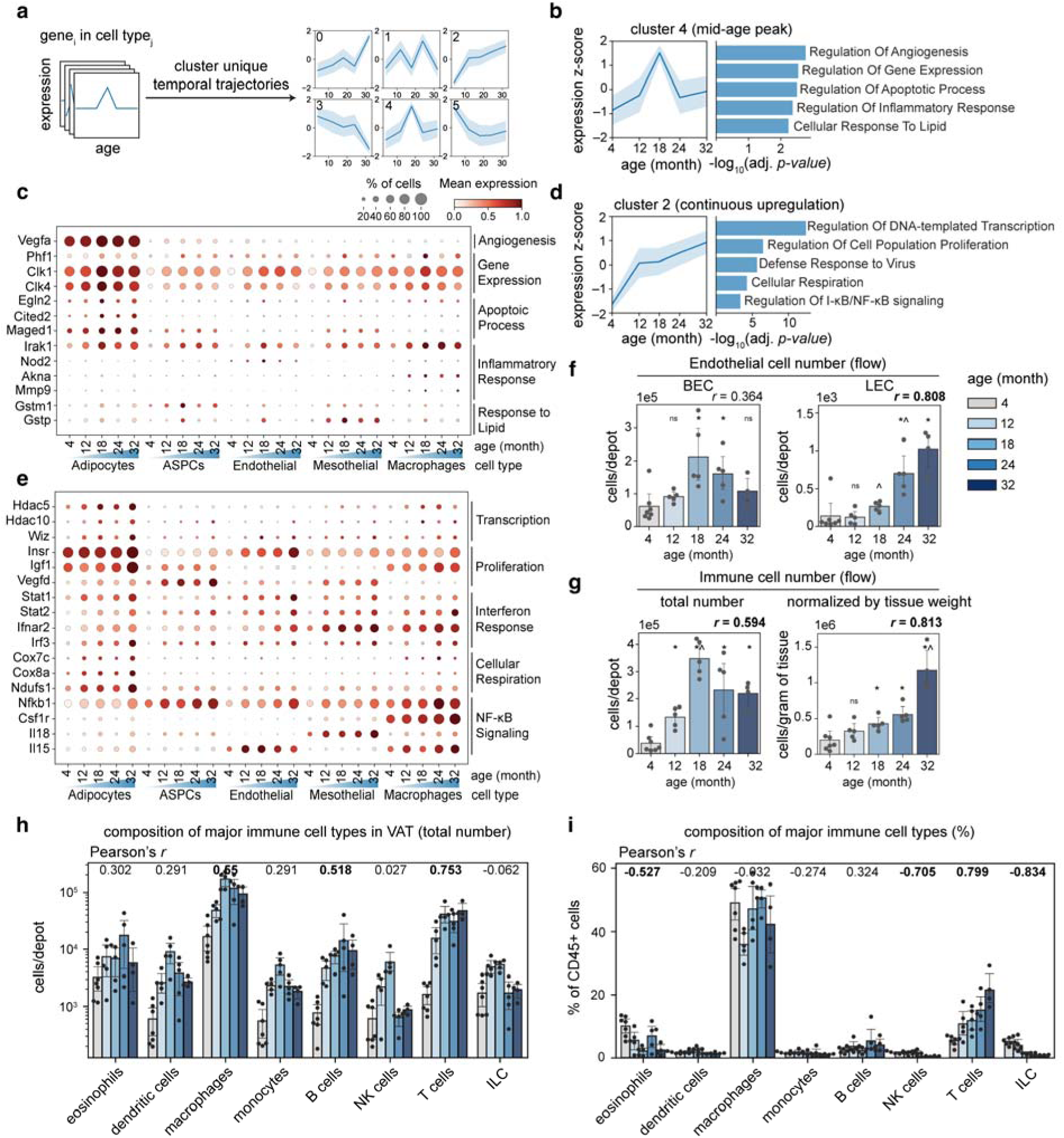
|. Adipose tissue experiences two distinct waves of inflammation during aging. **a**, Schematic for identifying distinct temporal expression trajectories among differentially expressed genes. **b**,**d**, Analysis of two key gene expression trajectories in VAT that correlated with the dynamics of tissue weight (**b**) and age (**d**). Average gene expression z-score (left) and significantly enriched Gene Ontology (GO) terms (right) are shown. **c**,**e**, Relative expression of genes identified in trajectories shown in (**b**) and (**d**), respectively. **f**, Absolute number of blood (BEC) and lymphatic (LEC) endothelial cells in VAT across all age groups (*n* = 4–7 per group). **g**, Total immune cell numbers (left) and weight-normalized immune cell density (right) in VAT across all age groups (*n* = 4–7 per group). **h**,**i**, Total number (**h**) and the relative proportion (**i**) of major immune cell types in VAT across all age groups. In each bar plot (**g**–**i**), data are presented as mean ± 95% confidence intervals, with each dot representing an individual mouse. Pearson’s correlation coefficients (r) with age are shown; bolded values indicate significant correlations (*p*-value < 0.05).

Among these trajectories, two specifically captured the dynamics of tissue weight and aging, respectively. The first trajectory mirrored the 18-month adipose tissue weight peak and was enriched for Gene Ontology (GO) terms characteristic of adipose tissue expansion and metabolic stress, such as angiogenesis (e.g., *Vegfa*) and response to lipid (e.g., *Gstm1*, *Gstp1*) (Fig. 3b,c). In contrast, the second trajectory exhibited a continuous increase with age and was enriched for GO terms associated with classic hallmarks of aging, including chromatin remodeling (e.g., *Hdac5*, *Hdac10*, *Wiz*) and mitochondrial dysfunction (e.g., *Cox7c*, *Cox8a*, *Ndufs1*) (Fig. 3d,e). Interestingly, both trajectories were enriched for unique inflammation-related GO terms, suggesting that adipose tissue experiences two distinct waves of inflammation. The 18-month peak trajectory included the innate immune signal transducers *Irak1* and *Nod2*, which mediate Toll-like and NOD-like receptor signaling, respectively (Fig. 3c). In contrast, the age-related trajectory was marked by a different inflammatory signature including interferon response (e.g., *Stat1*, *Stat2*) and NF-κB signaling (e.g., *Nfkb1*, *Il18*, *Il15*) (Fig. 3e).

We next sought to determine whether these transcriptomic signatures could predict shifts in the underlying cellular composition of the tissue. As a validation, we leveraged the distinct expression trajectories of *Vegfa* (peak at 18 month) and *Vegfd* (continuously upregulated), which are established regulators of blood vessel and lymphatic vessel growth, respectively (Extended Data Fig. 3c)^37^. Using flow cytometry, we confirmed that the total number of blood endothelial cells followed a trajectory similar to the expression of *Vegfa*, reaching a maximum at 18 months (Fig. 3c,f). As expected, *Vegfa* expression exhibited an inverse relationship with vascular density (Extended Data Fig. 3d–g), consistent with the known role of mild hypoxia in upregulating *Vegfa* as the tissue expanded^38^. Additionally, the number of lymphatic endothelial cells increased continuously with age, matching the expression trajectory of *Vegfd* (Fig. 3e,f). These findings suggested that these transcriptomic changes could reflect actual changes in cellular composition occurring within the aging adipose tissue.

We next used flow cytometry to identify the specific immune populations associated with each inflammatory wave (Extended Data Fig. 4a and Supplementary Table 4). We observed that although the total number of immune cells in adipose tissue followed a biphasic trajectory, it decreased only marginally between 18 and 32 months (Fig. 3g, left), in stark contrast to the substantial decrease in tissue weight (Fig. 1b). Consequently, the overall immune cell density increased continuously throughout the murine lifespan (Fig. 3g, right).

Among CD45^+^ immune cells, different cell types exhibited distinct temporal dynamics (Fig. 3h). The total number of dendritic cells, macrophages, monocytes, natural killer (NK) cells and innate lymphoid cells (ILCs) reached a maximum at 18 months before declining, aligning with the first, obesity-associated wave of inflammation. In contrast, T cell numbers remained elevated in older mice, resulting in a more pronounced increase in their density and an increased relative proportion among all immune cells (Fig. 3i and Extended Data Fig. 4b). This persistent accumulation was particularly notable because T cells are major producers of IFNγ^39^, and the IFNγ response was a key transcriptional signature enriched during the age-associated wave of inflammation. Within the T cell compartment, γδ T cells, effector helper T cells (CD4^+^, CD62L^-^, CD44^+^), and central memory cytotoxic T cells (CD8^+^, CD62L^+^, CD44^+^) exhibited the most significant age-related increases (Extended Data Fig. 4c–e).

Together, these findings indicated that VAT experiences two temporally, transcriptionally and cellularly distinct waves of inflammation during the natural aging process. Although the role of obesity in driving adipose tissue inflammation is well established^40^, the mechanisms by which aging independently contributes to adipose tissue inflammation remain less well understood. We therefore examined this less-explored, age-associated wave of inflammation in greater detail.

### Senescent cells induce an IFN**γ** response in adipose tissue

We first sought to identify potential drivers of this age-related IFNγ response signature. While the number of adipocytes generally increases as mice age, we found that even after correcting for this increase in cell number, the IFNγ response remained high in older tissues (Extended Data Fig. 5a,b). Interestingly, after correcting for age, we observed that this response was lower in samples with higher adipocyte number, suggesting a potential overlap between the driver of the age-associated IFNγ response and a mechanism that limits the generation of new adipocytes. Because cellular senescence is a hallmark process known to limit cell proliferation in both cell-autonomous and non-cell-autonomous fashion while simultaneously promoting inflammation through the secretion of pro-inflammatory factors^28,41–43^, we hypothesized it might be a driver of this response. Consistent with this hypothesis, we found that the proportion of cells expressing the senescence markers *p16* or *p21* increased with age, specifically within adipocytes, ASPCs and mesothelial cells (Fig. 4a,b and Extended Data Fig. 5c,d). Furthermore, the IFNγ response score correlated more strongly with the abundance of these *p16^+^* and *p21^+^* cells than it correlated with chronological age alone (Fig. 4c,d).

**Fig. 4.**
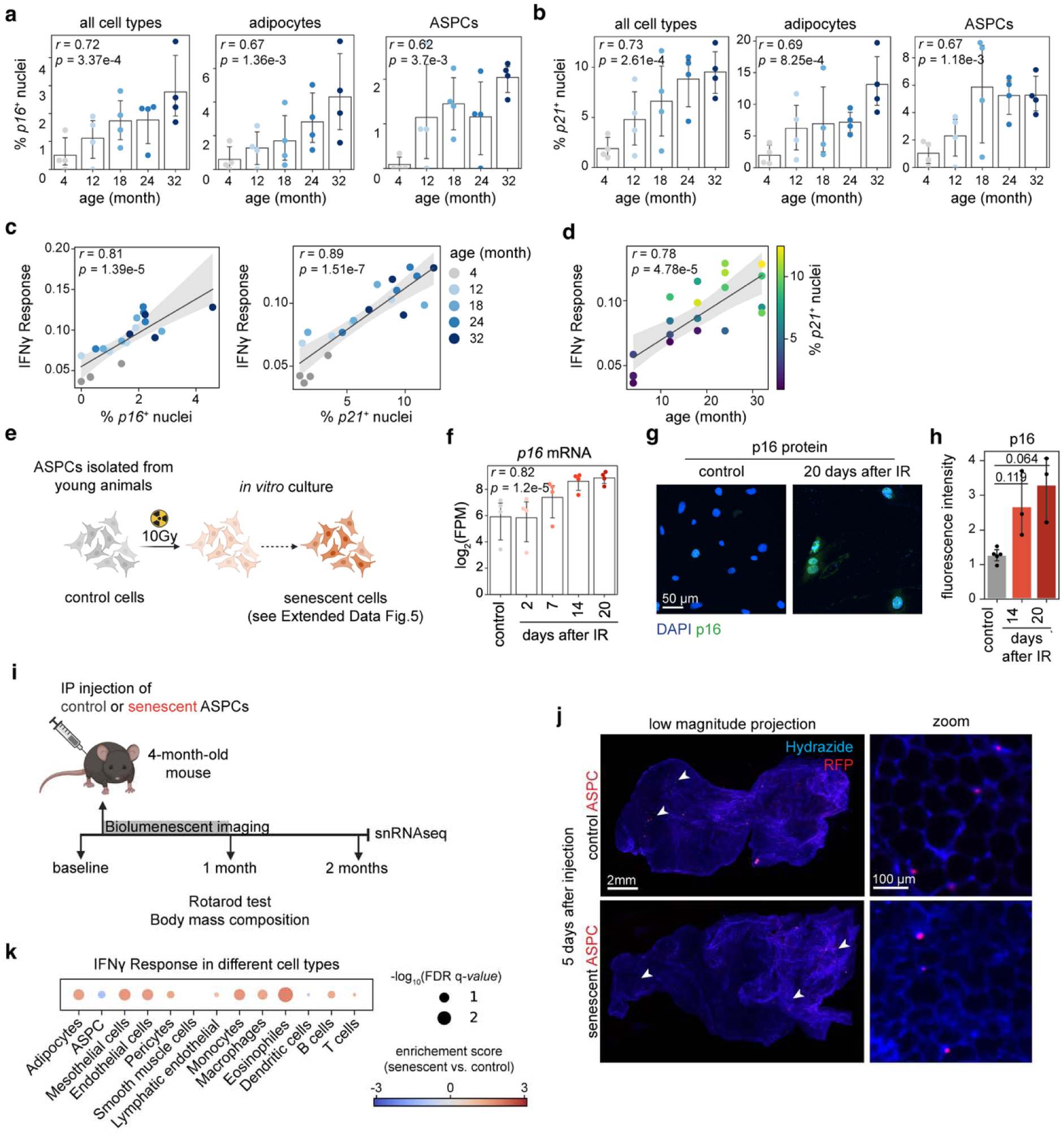
|. Senescent cells are sufficient to drive a subset of age-related changes in young VAT. **a**,**b**, Percentage of nuclei expressing *p16* (**a**) and *p21* (**b**) transcripts in all sequenced nuclei, adipocytes or ASPCs with age (*n* = 4 mice per group). **c**,**d**, Correlation between the percentage of *p16^+^* or *p21^+^* nuclei (**c**) or age (**d**) with the overall IFNγ response score of the tissue. **e**, Schematic showing the *in vitro* induction of cellular senescence in primary ASPCs. **f**, Expression of *p16* over time in ASPCs following irradiation (*n* = 4 per timepoint). **g**,**h**, representative immunofluorescence images (**g**) and quantification (**h**) of p16 protein levels in control versus senescent ASPCs (*n* = 3–5 per timepoint), *p*-values were determined by Welch’s two-sided t-test. **i**, Schematic of the senescent cell transplantation experiment. **j**, Representative images of entire VAT depots from mice that received control or senescent RFP^+^ ASPCs five days prior. We examined every organ within the peritoneal cavity and only observed RFP signal in VAT. **k**, GSEA result showing the effect of senescent cells on the IFNγ response score across various cell types in VAT compared to controls. In each bar plot (**a**, **b**, **f** and **h**), data are presented as mean ± 95% confidence intervals, with each dot representing an individual mouse or an independent biological replicate (cells isolated from different mice). For plots in (**a**, **b**, **c**, **d** and **f**), Pearson’s correlation coefficient (*r*) and *p*-value are shown.

To move beyond correlation and experimentally test the causal role of senescent cells in initiating the age-associated IFNγ response, we sought to determine whether their presence was sufficient to induce this signature in young mice. We utilized ASPCs for this experiment because they represented a key senescent population identified in our aging data and were amenable to *ex vivo* manipulation and transplantation. Using irradiation as a canonical senescence inducer, we confirmed the induction of a stable senescent state in primary ASPCs, as evidenced by cell cycle arrest (Extended Data Fig. 5e–g), the induction of p16 and p21 (Fig. 4f–h and Extended Data Fig. 5h), downregulation of lamin B1 (Extended Data Fig. 5i–k) and the development of a senescence-associated secretory phenotype (SASP) comprising many inflammatory cytokines (Extended Data Fig. 5l)^44,45^. By verifying these hallmarks before transplantation, we ensured that any subsequent phenotypes in the recipient mice were driven by a *bona fide* senescent population.

We injected these senescent ASPCs into young mice intraperitoneally, following a previously established protocol (Fig. 4i and Supplementary Table 5)^46^. We confirmed that both control and senescent ASPCs accumulated specifically in VAT (Fig. 4j) and were cleared within two weeks (Extended Data Fig. 6a). Consistent with the prior report, this transient exposure to senescent cells was sufficient to induce physical dysfunction in young mice, as evidenced by impaired rotarod performance (Extended Data Fig. 6b), notably occurring without altering body mass composition or glucose tolerance (Extended Data Fig. 6c,d).

To investigate the impact of senescent cells on gene expression, we performed single-nucleus RNA sequencing on VAT of recipient mice. For this analysis, we utilized a droplet-based platform to maximize nuclei recovery, particularly for immune cell populations with inherently lower mRNA content. This approach allowed us to capture a broader range of immune cells compared to the combinatorial barcoding-based approach (Extended Data Fig. 7a). While this short exposure to senescent cells did not recapitulate all age-dependent transcriptional changes, it induced an IFNγ response across many cell types (Fig. 4k, Extended Data Fig. 7b and Supplementary Table 6). Notably, this induction occurred without an associated change in adipocyte size (Extended Data Fig. 7c), suggesting that an increased IFNγ response is likely not the primary driver of the reduction in adipocyte size observed in late life. Furthermore, total adipocyte number remained unchanged (Extended Data Fig. 7c). While we hypothesized that senescent cells could simultaneously drive the IFNγ response and limit the production of new adipocytes in aged animals, the lack of change in adipocyte number in these young recipients is likely because hyperplastic expansion of the VAT initiated only after 12 months of age (Fig. 1f).

In addition to the core IFNγ response signature, we also observed upregulation of complement system components, a feature that also occurs in VAT with age (Extended Data Fig. 7d). Together, these results demonstrated that a short exposure to senescent cells is sufficient to drive an IFNγ response in young VAT, supporting a causal relationship between cellular senescence and the age-associated inflammatory wave.

### IFN**γ** response induced by senescent cells upregulates MHC II in the vasculature

Among the transcriptional changes induced by senescent-cell transplantation, we observed an unexpected upregulation of major histocompatibility complex (MHC) class II molecules in endothelial cells (Fig. 5a). This induction was also observed in the vasculature of naturally aged adipose tissues (Fig. 5a). Expression of MHC II is typically restricted to professional antigen-presenting cells (APCs), where it presents antigens to CD4^+^ T cells for their activation (Fig. 5b). We also observed upregulation of the invariant chain *Cd74* and baseline expression of the peptide-loading machinery *H2-DMa* and *H2-DMb1* (Fig. 5a,b)^47^, suggesting the molecular machinery required for antigen processing and presentation was present.

**Fig. 5.**
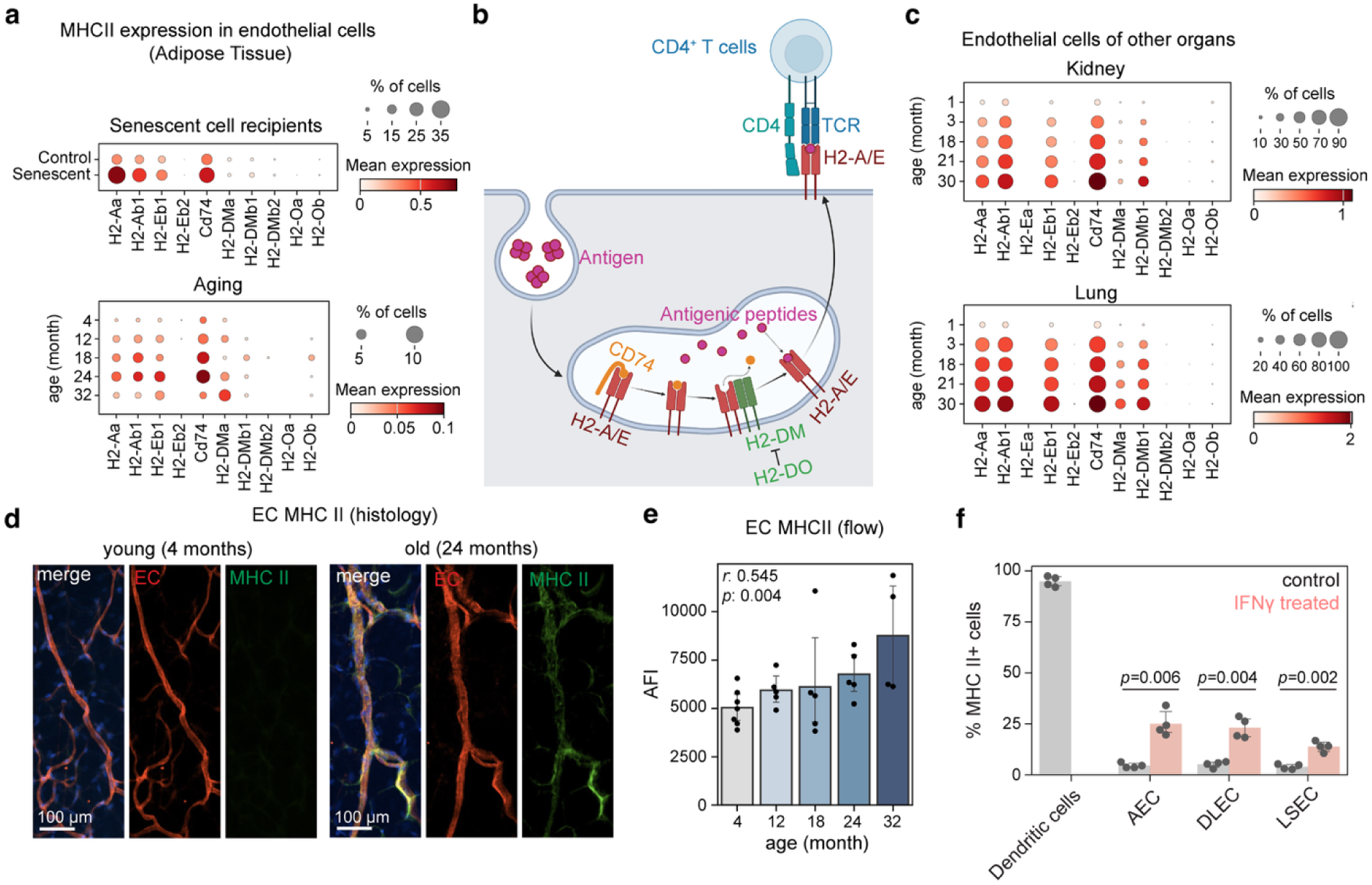
|. Senescent cells and aging upregulate MHC II on the vasculature. **a**, Expression of MHC class II-associated components in VAT endothelial cells following senescent cell transplantation (top) and across the murine lifespan (bottom). **b**, Schematic illustrating antigen presentation by MHC II. **c**, Expression of MHC II molecules in endothelial cells of kidney and lung across different ages. **d**, **e**, MHC II protein expression on VAT endothelial cells (EC) of young and aged mice by immunofluorescent staining (**d**) and flow cytometry (**e**). **f**, Quantification of the percentage of primary endothelial cells expressing MHC II with or without IFNγ treatment.

While endothelial MHC II expression has been reported in specific disease contexts and on specialized endothelia^48,49^, its appearance in aging was only recently noted in the brain^50^. To determine if this was a general feature of aging, we analyzed the Tabula Muris Senis single-cell RNAseq dataset^28^ and found that age-associated endothelial MHC II upregulation is a widespread phenomenon, occurring across numerous organs, including the kidney, lung, pancreas and heart (Fig. 5c and Extended Data Fig. 7e). We confirmed this upregulation at the protein level in VAT using immunofluorescent staining and flow cytometry (Fig. 5d,e).

Given that the IFNγ response increased with age in adipose tissue and IFNγ is a potent inducer of MHC II ^51,52^, we hypothesized that elevated IFNγ levels drive MHC II expression on endothelial cells. Indeed, treating primary mouse endothelial cells with IFNγ robustly induced MHC II *in vitro* (Fig. 5f and Extended Data Fig. 8a). The expression of MHC II was lower in endothelial cells compared to dendritic cells, but the expression persisted for at least three days following IFNγ removal (Extended Data Fig. 8a). We tested whether conditioned media from senescent ASPCs induced MHC II expression (Extended Data Fig. 8b), and found that it did not, consistent with our transcriptomic data showing that senescent cells themselves did not express *Ifng* (Extended Data Fig. 8c). This supported a model where senescent cells initiated the IFNγ response through an intermediate cell type, which in turn upregulated MHC II on the vasculature.

### Vascular MHC II facilitates trans-endothelial extravasation of CD4^+^ T cells

Given that MHC II interacts with the T cell receptor on CD4^+^ T cells, we asked if there was a corresponding change in CD4^+^ T cells in response to senescent cell transplantation. Flow cytometry analysis revealed that the transplantation of senescent cells led to a specific and significant increase in effector CD4^+^ T cell number in VAT (Fig. 6a, Extended Data Fig. 9a–c and Supplementary Table 7), mirroring the accumulation observed in naturally aged VAT (Fig. 6a). Consistent with this finding, the clearance of senescence cells using senolytics has been shown to reduce the abundance of T cells in aged VAT^53^. Furthermore, analysis of the Tabula Muris Senis bulk RNAseq dataset^22^ showed increased *Cd4* transcripts with age across many of the same organs exhibiting vascular MHC II upregulation, suggesting a corresponding accumulation of CD4^+^ T cells in these organs as well (Fig. 6b).

**Fig. 6.**
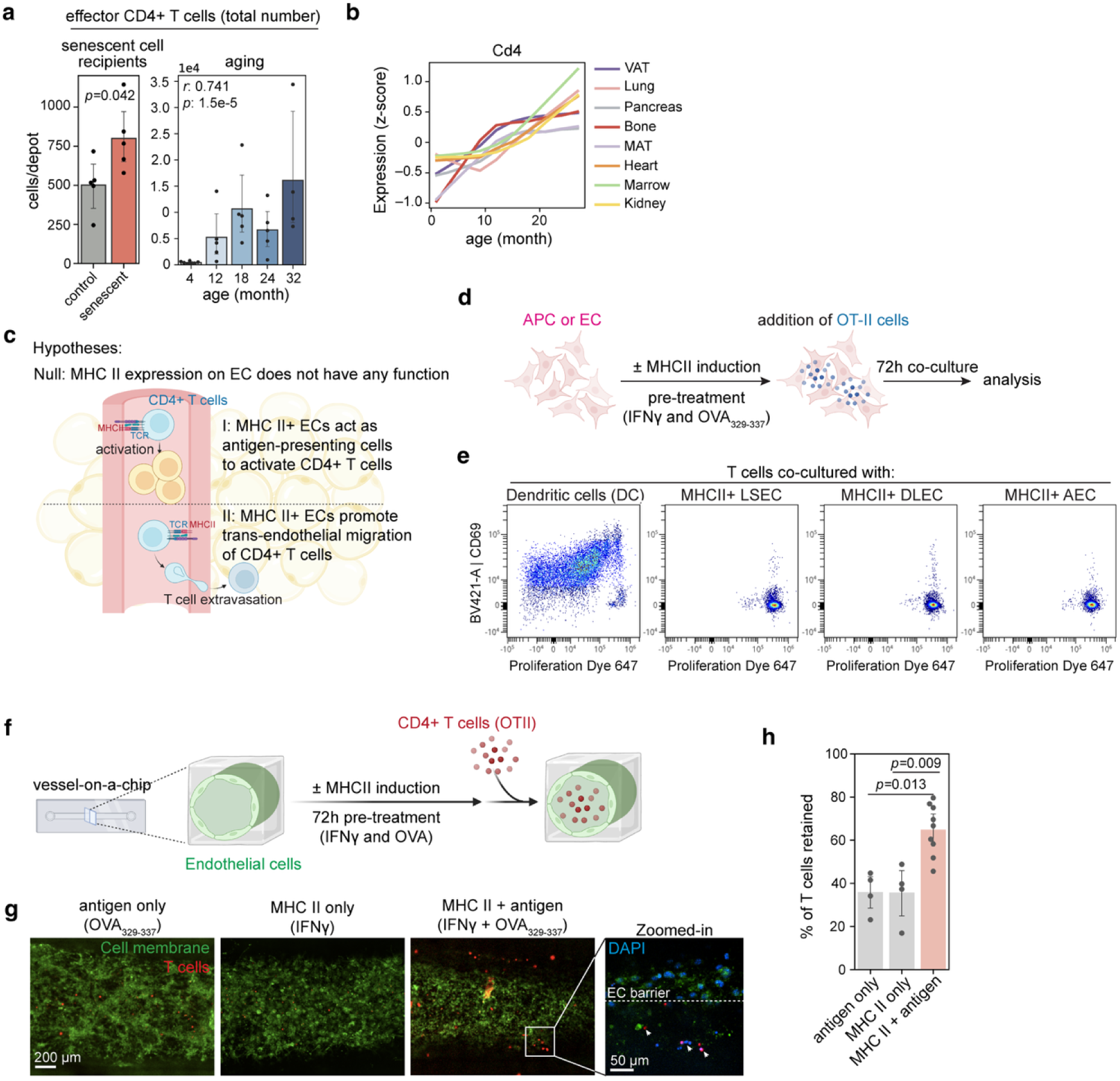
|. Vascular MHC II facilitates trans-endothelial migration of CD4^+^ T cells. **a**, Number of activated CD4^+^ T cells in VAT following senescent cell transplantation (left) and across the murine lifespan (right) (n=4–7 per group). **b**, Relative expression of *Cd4* in different tissues with age. **c**, Proposed hypotheses for the function of vascular MHC II. **d**, Schematic of the *in vitro* T-cell activation assay. **e**, Activation and proliferation status of naïve CD4^+^ T cells after co-culturing with dendritic cells (positive control) or MHC II-expressing endothelial cells. **f**, Schematic of the vessel-on-a-chip microfluidic platform used to model trans-endothelial migration. **g**,**h**, Representative confocal images (**g**) and quantification (**h**) of CD4^+^ T-cell retention and migration through endothelial vessels. In each bar plot (**a** and **h**), data are presented as mean ± 95% confidence intervals, with each dot representing an individual mouse or an independent vessel-on-a-chip device. Statistical significance was determined by Welch’s two-sided t-test or Pearson’s correlation.

These data raised the possibility of a functional link between vascular MHC II and the accumulation of CD4^+^ T cells. To test the functional consequences of vascular MHC II on CD4^+^ T cells, we formally evaluated two hypotheses: first, MHC II enables endothelial cells to activate naïve CD4^+^ T cells, akin to professional APCs; or second, MHC II facilitates the extravasation of CD4^+^ T cells from the circulation into the tissue (Fig. 6c).

To test the first hypothesis, we co-cultured naïve CD4^+^ OT-II T cells with MHC II-expressing endothelial cells pre-incubated with their cognate antigen peptide, OVA_329-337_ (Fig. 6d). Unlike dendritic cells, which served as a positive control, endothelial cells failed to induce T cell activation or proliferation (Fig. 6e and Extended Data Fig. 8d), even though two out of the three endothelial cell lines we tested also expressed the co-stimulatory molecule CD80 (Extended Data Fig. 8e). This result suggested that unlike classical APCs, endothelial cells likely could not activate naïve T cells.

We then tested the second hypothesis using a vessel-on-a-chip model to visualize T cell migration directly (Fig. 6f). We observed that vessels expressing MHC II with the cognate antigen peptide retained significantly more CD4^+^ OT-II T cells than did control, non-MHC II-expressing vessels (Fig. 6g,h). Moreover, we observed clear extravasation of T cells across the endothelial layer exclusively in the MHC II-expressing vessels (Fig. 6g). Importantly, this effect was antigen-dependent, as IFNγ treatment without the cognate antigen failed to promote retention and trans-endothelial migration of T cells (Fig. 6g,h).

Together, these results supported the hypothesis that MHC II expression on endothelial cells facilitated the antigen-dependent extravasation of cognate CD4^+^ T cells, suggesting a possible mechanistic explanation for the age-dependent accumulation of CD4^+^ T cells in adipose tissue.

### CD4^+^ T cells are required for the age-related IFNγ response and immune cell accumulation

Our findings thus far suggested that senescent cells indirectly promoted the IFNγ response that upregulated MHC II in the vasculature, which in turn facilitated the extravasation of CD4^+^ T cells into surrounding tissues. To identify cellular sources of IFNγ, we analyzed the cytokine-producing capacity of the immune compartment within the VAT and found that CD4^+^ T cells represented a predominant source of IFNγ in aged VAT (Fig. 7a). Thus, CD4^+^ T cells might provide the very signal required to maintain their own recruitment and sustain the chronic inflammation observed during aging.

**Fig. 7.**
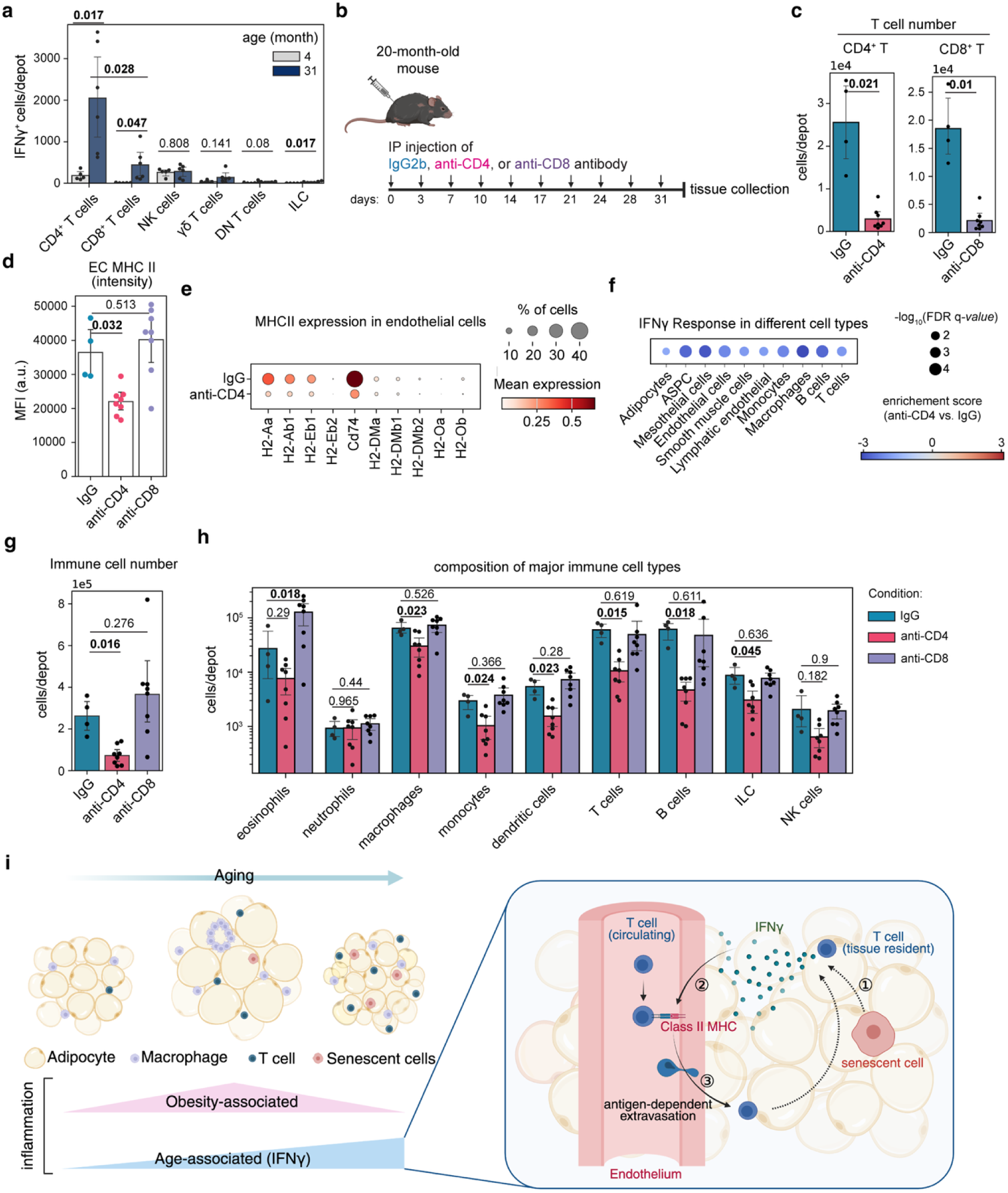
|. CD4^+^ T cells are required for the age-related immune cell accumulation and IFNγ response. **a**, Absolute number of IFNγ^+^ cells quantified by flow cytometry, stratified by cell type within the VAT of young versus old mice. **b**, Schematic of the *in vivo* T-cell depletion experiment in aged mice. **c**, Number of CD4^+^ or CD8^+^ T cells in VAT following treatment with depleting antibodies or isotype controls. **d**,**e**, Expression of MHC II on VAT endothelial cells (CD45^-^ CD31^+^) measured by flow cytometry (**d**) and single-nucleus RNA sequencing (**e**). **f**, GSEA result showing the effect of CD4^+^ T-cell depletion on the expression of IFNγ response genes across various cell types in aged VAT compared to isotype IgG control, **g**,**h**, Absolute numbers of total immune cells (**g**) and major immune subtypes (**h**) in aged VAT following T-cell depletion. **i**, Model: a positive feedback loop that sustains age-associated inflammation. Senescent cells initiate the loop by indirectly promoting an initial IFNγ response that upregulates vascular MHC II. Vascular MHC II promotes the recruitment of more CD4^+^ T cells into the tissue, which in turn produce more IFNγ. Data are presented as mean ± 95% confidence intervals, with each dot representing an individual mouse (n=4–8 per group). Statistical significance was determined by Welch’s two-sided t-test.

To test this hypothesis, we depleted CD4^+^ T cells from 20-month-old female mice (Fig. 7b). We also included a CD8^+^ T-cell depletion group as a control, as these cells also accumulated in VAT with age and produced IFNγ but were not implicated in the MHC II-mediated recruitment pathway (Extended Data Fig. 4c and Fig. 7a). Following a one-month depletion regimen, we confirmed the successful removal of each T-cell subset in the VAT (Fig. 7c, Extended Data Fig. 10d and Supplementary Table 8), providing a robust model to assess their relative contribution to the age-associated inflammation.

We observed that depletion of CD4^+^ T cells, but not CD8^+^ T cells, led to a significant decrease in MHC II expression on endothelial cells (Fig. 7d,e). This finding provided *in vivo* support for the hypothesis that CD4^+^ T cells are required for the age-related upregulation of vascular MHC II. Consistent with this reversion of the activated endothelial state, CD4^+^ T-cell depletion also significantly reduced expression of IFNγ response genes across multiple cell types in aged VAT (Fig. 7f, Extended Data Fig. 10a and Supplementary Table 9).

Strikingly, the depletion of CD4^+^ T cells, but not CD8^+^ T cells, also led to a dramatic reduction in the total number of immune cells in aged VAT (Fig. 7g), reverting the inflammatory burden to levels lower than 12-month-old mice (Fig. 3g). This reduction was not simply due to loss of T cells, as we observed a broad normalization of the inflammatory infiltrate, including significant reductions in the number of macrophages, monocytes, dendritic cells, B cells and ILCs (Fig. 7h and Extended Data Fig. 10e). Importantly, this effect was specific to the aged tissue and was observed in both sexes, as CD4^+^ T-cell depletion in old but not young male mice showed an reduction in total immune cell number and specific immune cell subtypes (Extended Data Fig. 10d,e). Furthermore, these inflammatory shifts occurred without changes in tissue weight, adipocyte size, or adipocyte number (Extended Data Fig. 10f–h).

In summary, our results demonstrated that CD4^+^ T cells are required for the sustained age-related IFNγ response and immune cell accumulation in aged adipose tissue. They support a model in which CD4^+^ T cells play a central role in a positive feedback loop initiated by senescent cells—IFNγ produced by CD4^+^ T cells upregulate vascular MHC II to promote tissue infiltration of additional CD4^+^ T cells, which in turn sustain the chronic inflammatory state of aging (Fig. 7i).

## Discussion

In this study, we provided a multi-modal atlas of aging adipose tissue that decoupled the correlates of chronological aging from those of mid-life adiposity (Fig. 1 and 2). By examining both gene expression and immune cell composition across the murine lifespan, we demonstrated that the visceral adipose tissue experienced two distinct waves of inflammation: a mid-life wave of obesity-associated inflammation and an age-specific wave characterized by a heightened IFNγ response (Fig. 3). Together, our data suggest a molecular basis for this age-associated wave that incorporates a positive inflammatory feedback loop. In this model, senescent cells trigger an initial IFNγ signal (Fig. 4) that upregulates vascular MHC II (Fig. 5), thereby facilitating the movement of CD4^+^ T cells from the circulation into the tissue (Fig. 6). Once recruited, these CD4^+^ T cells produce more IFNγ, thus perpetuating and enhancing the cycle of T cell accumulation and chronic inflammation (Fig. 7).

Our observation that the first wave of inflammation tracked with the peak in adipose tissue weight identified adiposity as a critical confounding factor when interpreting changes in adipose tissue during aging. Because *ad libitum*-fed mice often exhibit significantly increased fat weight at middle age^18–20^, the structural and molecular changes observed during this period are likely conflated by the impacts of both obesity and chronological aging. Our multi-age strategy allowed us to distinguish these factors, revealing that the two waves of inflammation involved distinct sets of genes and immune cell populations.

Although we focused on the age-associated inflammatory wave in the current study, it is important to note that mid-life obesity may contribute to the development of the age-associated wave. For instance, obesity is a known inducer of cellular senescence in adipose tissue^54,55^, which could in turn accelerate and amplify the age-associated IFNγ response. It would therefore be informative for future studies to test whether these age-associated inflammatory signatures can be delayed or ameliorated in mice that do not experience significant mid-life weight gain.

These investigations could clarify the extent to which maintaining metabolic homeostasis influences the timing and intensity of age-associated inflammation.

Our investigation into the age-associated inflammatory wave led to the discovery of an age-dependent upregulation of MHC II on the vasculature. While vascular MHC II expression has been described previously in specific diseases and specialized tissue types^56–59^, our findings revealed its role as a regulator of CD4^+^ T cell extravasation during aging. Furthermore, the antigen-dependency observed in our vessel-on-a-chip model suggested that the recruitment of CD4^+^ T cells might be tuned to specific signals presented on the aging vasculature. This is in line with a report in a multiple sclerosis model, in which the vasculature presents myelin peptide to promote the trans-endothelial migration of myelin-reactive T cells^60^. It is interesting to speculate that during aging, endothelial cells could take up antigens from surrounding tissue parenchyma and then present them on luminal MHC II complexes to circulating CD4^+^ T cells. Although the identity of antigens presented on the aging vasculature remains a critical question for future inquiry, such a mechanism implies that aging remodels the vasculature into an active immunological interface. This mechanism can operate alongside the progressive loss of vascular integrity and the upregulation of classic immune cell recruitment signals for the recruitment and accumulation of immune cells in aged tissues^61–63^.

This selective recruitment of CD4^+^ T cells mediated by vascular MHC II suggests an ordered sequence of events that drives age-associated inflammation. While senescent cells have long been recognized as a driver of age-related inflammation through the SASP, our findings suggest a specific role for senescent cells as upstream initiators of a CD4^+^ T cell-mediated positive feedback loop. By establishing this relay, even a small number of senescent cells could potentially initiate an inflammatory state that is subsequently amplified and maintained by CD4^+^ T cells. This model provides an explanation for how a small number of senescent cells found in aged tissues can exert a disproportionate, tissue-wide effect.

In summary, this work provides new insights into how age-related inflammation is initiated and sustained by identifying a self-sustaining positive feedback loop that links cellular senescence, vascular MHC II and immune cell infiltration. Our work raises many exciting questions for future studies, including identifying the particular SASP components responsible for triggering the initial IFNγ response, characterizing the antigen(s) presented by MHC II on the aging vasculature, validating the generalizability of this mechanism across species, and exploring the therapeutic potentials of disrupting this cycle to ameliorate age-related inflammation and diseases.

## Acknowledgments

We thank Calico Genomics Group (Wenjun Kong, Po-Han Tai, Evangelia Malahias, Amy Jo Johnson, Ikenna Anigbogu, Andrew Keyser, and David Hendrickson), Calico Histology and Spatial Biology group (Patrick Godfrey, Aaron Stebbins, Varahram Shahryari and James Lee), Calico Laboratory Animal Resources (Samira Nazertehrani, Natalie Smolnikov, Raul Garcia-Gonzalez and Ellie Karlsson), Calico Physiology Lab (José Zavala-Solorio and Chunlian Zhang), Calico Flow Cytometry Lab (Ka Man Li and Andrew Nguyen) for guidance and assistance with experiments. We thank Michael Lenardo, Fiona Harding, Oliver Hahn, Seonhui Shim, Thao Nguyen, Wen Lu, Yi Ding and members of the Kenyon Lab for insightful feedback on this study. All schematic illustrations were created using BioRender.

## Author contributions

Q.X., and C.K. conceived the project. Q.X. performed all experiments and analyses, with contributions from other authors indicated below. T.D.L., S.J., A.D.F., M.P., and H.T.A. assisted with tissue collection and sample processing with input from C.J.. A.S. generated vessel-on-a-chip devices and performed data analysis with input from K.N.. K.H. imaged whole-mount adipose tissues. AEYTL performed vessel segmentation and assisted with histology image analysis. M.J. performed metabolic modeling. Q.X., and C.K. wrote the manuscript, and all authors provided feedback.

## Competing interests

All authors are employees of Calico Life Sciences, LLC at the time of this study.

## Methods

### Animals

All animal experiments were conducted under protocols approved by the Institutional Animal Care and Use Committee (IACUC) at Calico Life Sciences LLC. C57BL/6J (Jax #000664), *CAG-mRFP1/J* (Jax #005884)^64^, OT-II (Jax #004194)^65^, and *Actin-Cre* (Jax #033984)^66^; *Rosa26^LSL-OVA-Luc^*^67^ mice were purchased from the Jackson Laboratory. All mice were acclimated for a minimum of one week in the Calico animal facility before experiments commenced. Mice were housed under a 12-hour light-dark cycle with *ad libitum* access to standard chow diet and water, unless otherwise indicated.

### Tissue collection

Tissues were collected between 8:00 AM and 12:00 PM to minimize circadian variation. Mice were deeply anesthetized with isoflurane in a clean chamber, weighed, and then euthanized by cardiac exsanguination. For flow cytometry analysis, mice were transcardially perfused with 15 ml of PBS prior to tissue collection. Subcutaneous adipose tissue was collected from the inguinal region. Inguinal lymph nodes were carefully removed from SAT intended for single-nucleus RNA sequencing or flow cytometry experiments. Visceral adipose tissue was collected from the perigonadal region. For paired histological analysis and transcriptome profiling, one side of the adipose tissue was fixed in 10% formalin, and the contralateral side was flash frozen and stored in liquid nitrogen for subsequent nuclei isolation.

### Histology and immunohistochemistry

Adipose tissues and livers were fixed overnight in 10% neutral buffered formalin. After fixation, tissues were rinsed with phosphate-buffered saline (PBS) and processed on an automated tissue processor (Tissue-Tek VIP6 AI, Sakura) following a standard dehydration and clearing protocol. Samples were then infiltrated with molten paraffin wax (Paraplast, Leica Biosystems) at 60 °C for 3 hours and embedded into paraffin blocks using a tissue embedding station (HistoCore Arcadia H, Leica). Paraffin blocks were sectioned at 6 µm using a rotary microtome (HistoCore AUTOCUT, Leica). Immunohistochemistry was performed on the Bond RX Automated IHC Stainer (Leica Biosystems) using a rabbit anti-CD31 antibody (1:100, Cell Signaling Technology), ER2 antigen retrieval solution, and the Bond Polymer Refine Detection System (DS9800, Leica Biosystems), followed by hematoxylin counterstaining. Slides were mounted and scanned on a whole-slide scanner (Axioscan 7, Zeiss or VS200, Olympus). Tissue sectioning, H&E staining, and image acquisition were performed either in-house or by a commercial laboratory (Histowiz).

### Morphometric quantification

Image analysis for adipose and liver tissues was performed using a custom Python script. Briefly, images were downsampled by a factor of 4 and converted to a single grayscale channel by taking the minimum intensity across channels. A binary mask was created using a Gaussian-Laplace filter and refined through a sequence of morphological operations (binary opening, binary closing, and binary dilation). From the refined binary mask, internal voids within connected components (individual adipocytes or lipid vacuoles) were filled, and morphological features of each connected component were extracted using “skimage.measure.regionprops”.

For adipose tissue, objects that were too small (less than 226 µm^2^, likely noise), too large (larger than 15000 µm^2^, likely merged objects and non-adipocyte structures), or not sufficiently round (sphericity less than 0.35, likely non-adipocyte structures) were filtered out. Mean cross-sectional area (CSA) was calculated by averaging the size of all segmented adipocytes from each sample. Total adipocyte number for each sample was estimated using the following formula, adapted from a previously published method^68^:

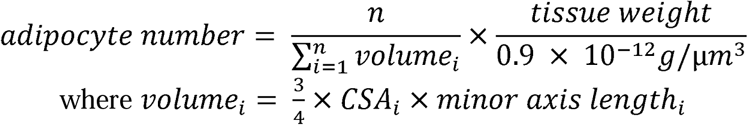

Here, n represents the total number of segmented adipocytes, and i refers to each segmented adipocyte.

For liver tissue, lipid vacuoles are typically smaller and more spherical compared to adipocytes in adipose tissue. Therefore, objects that were too large (larger than 4500 µm^2^), not round (sphericity less than 0.7), or too irregular (solidity less than 0.9) were filtered out. The percentage of lipid vacuole area was calculated using the following formula:

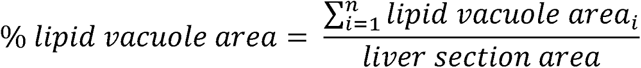

Here, n represents the total number of segmented lipid vacuoles, and i refers to each segmented lipid vacuole.

### Single-nucleus RNA sequencing

Nuclei were isolated from frozen adipose tissue samples using a modified protocol adapted from a previously described method ^31^. For each sample, approximately 50 mg of frozen adipose tissue was placed into a gentleMACS C tube (Miltenyi Biotec) containing 2 ml of freshly prepared TST buffer (10 mM Tris-HCl, 146 mM NaCl, 1 mM CaCl_2_, 21 mM MgCl_2_, 0.01% BSA, 0.03% Tween 20, and 0.2 unit/μL RNasin Plus). Tissues were dissociated using a gentleMACS Dissociator (Miltenyi Biotec) with the program “mr_adipose_01” performed twice, including a 5-minute incubation on ice between each run. After an additional 10-minute incubation on ice, the lysate was passed through a 40 µm nylon filter and collected into a 50 mL conical tube on ice. The filter was rinsed with 3 mL of freshly prepared ST buffer (10 mM Tris-HCl, 146 mM NaCl, 1 mM CaCl_2_, 21 mM MgCl_2_ and 0.2 unit/μL RNasin Plus). Samples were then centrifuged at 500 g for 5 minutes at 4 °C with the brake set to low. The nuclear pellet was washed three times by resuspending in resuspension buffer (1% BSA and 0.2 unit/μL RNasin Plus in PBS) followed by centrifugation. A 10 μL aliquot was used to determine nuclei quality and count using NucleoCounter NC-3000 (ChemoTetec) after staining with Solution 13 (AO•DAPI, ChemoTetec).

For sequencing the aging cohort using the split-pool combinatorial barcoding sequencing approach, nuclei were fixed according to the low volume fixation protocol (ScaleBio). The fixed samples were stored in -80°C until all samples were collected and fixed for loading (within 6 months of storage). The samples were processed using ScaleBio single-cell RNA sequencing kit (v1.1). Briefly, fixed samples were thawed on ice, recounted, and adjusted to 2,000 nuclei/ml with ScaleBio wash buffer (Part No. 202100001) prior to loading. Each sample was loaded into 2 wells of the initial RT 96-well plate with 10,000 nuclei per well. The plate was incubated, and nuclei in the wells were pooled to re-distribute into the second 384-well ligation plate. Following ligation, nuclei were pooled again and counted to distribute into the final distribution plates (96-well plates) with 1,600 nuclei per well. The library was prepared and pooled from the final distribution plates with a target recovery of 100,000 nuclei per final distribution plate. The final library was loaded at a concentration of 2.7nM.

For the droplet-based sequencing approach, nuclei were immediately loaded onto a 10x Chromium controller (10x Genomics) according to the manufacturer’s protocol. For each sample, 10,000 nuclei were loaded into one channel of a Chromium Chip (10x Genomics) and libraries were prepared following the Single Cell Gene Expression v3.1 (senescent cell transplantation experiment) or v4 (T cell depletion experiment) protocol.

Libraries were sequenced on a NovaSeq 6000 system (Illumina). Raw sequencing reads were demultiplexed to FASTQ format files using bcl2fastq (Illumina).

### Single-nucleus RNAseq read alignment and preprocessing

For reads generated using ScaleBio single-cell RNA sequencing kit, data was processed by the ScaleRNA pipeline (v1.5.0) and mm10 mouse reference genome was used to generate the gene-by-cell count matrices. For the droplet-based single-nucleus RNA sequencing data, reads were aligned using CellRanger (version 7.0.0) against the GRCm39 mouse reference genome.

CellBender (version 3.7) was applied to the ASPC recipient samples to remove ambient RNA from the count matrices^69^.

Subsequent filtering and preprocessing were performed using Scanpy (version 1.9.3)^70^. Nuclei with less than 1,000 unique molecular identifiers (UMI), fewer than 200 detected genes, or identified as probable doublets by DoubletDetection were filtered out. The filtered data were then normalized using a scaling factor of 10,000 and log-transformed.

### Dimensionality reduction, clustering, and annotation

We largely followed standard procedures for variable gene selection, dimensionality reduction, and clustering using Scanpy. Briefly, Principal Component Analysis (PCA) was performed on highly variable genes, and the top 20 principal components were used for UMAP dimensionality reduction. Leiden clustering was applied to define distinct clusters.

Clusters were manually annotated using previously reported marker genes^31,71^. Clusters exhibiting high *Ptprc* expression were isolated and re-clustered to further resolve individual immune cell types. To confirm the cell type annotation, automatic cell type prediction was performed using over-representation analysis implemented in decoupler (version 1.5.0)^72^ against the PanglaoDB database^73^.

### Differential expression analysis between SAT and VAT

To understand the transcriptional differences between SAT and VAT (Fig. 2c), we performed differential gene expression analysis using a pseudobulk approach. Briefly, raw counts from the same samples were summed across cell types. For each age, pyDESeq2 was applied to model the count data with a negative binomial distribution, accounting for size factors and dispersion estimates^74,75^. A contrast on depot type was applied to identify gene expression differences between SAT and VAT. Following differential gene expression analysis, Gene Set Enrichment Analysis (GSEA) was performed to identify biological pathways significantly enriched in each depot at each age. Genes were ranked based on their Wald statistic from the DESeq2 results, and enrichments of the MSigDB Hallmark gene sets were determined using the “get_gsea_df” function in decoupler (version 1.5.0)^72^.

### Metabolic modeling

To infer the metabolic state of adipocytes, pseudo-bulk expression profiles of adipocytes were generated using a bootstrapping approach. To account for the potential impact of cell cycle on gene expression, a random sample of 50% of nuclei within each predicted cell cycle phase (G1, S, and G2/M) was drawn from each sample, and raw counts were summed to create a pseudo-bulk profile. This process was repeated 10 times for each sample to create a robust set of pseudo-bulk profiles for metabolic modeling analysis by the Compass algorithm^36^. The differential potential activity of each reaction was assessed with Cohen’s d statistic, and *p-values* were calculated using the Wilcoxon rank-sum test.

### Temporal gene expression trajectory analysis

Pseudo-bulk expression profiles were generated by summing the raw counts of each cell type within each sample using the decoupler-py package^72^. We focused on cell types that had a substantial number of nuclei sequenced across all five age points, including adipocytes, ASPCs, macrophages, endothelial cells, and mesothelium cells.

For each selected cell type, differential expression analysis was performed on these pseudo-bulk profiles across every pair of ages using pyDESeq2. Genes with an adjusted *p-value* of less than 0.05 in at least one pairwise age comparison were considered differentially expressed for that specific cell type. Subsequently, the expression profiles of these cell-type-specific differentially expressed genes were determined. This involved calculating the average expression of each gene at each age, followed by standardizing the expression values using “sklearn.preprocess.StandardScaler” to center the data and scale it to unit variance.

The scaled expression profiles from all analyzed cell types were then combined. PCA was performed to transform the combined expression profiles into a lower-dimensional space. K-means clustering was applied to the PCA-transformed data using “sklearn.cluster.KMeans”. The optimal number of clusters was determined by evaluating a range of n_clusters parameters. A value of 6 was chosen as it captured all significant temporal patterns while avoiding redundant clusters. Over-representation analysis of gene ontology terms was performed on the combined gene set using the enrichR package.

### Flow cytometry analysis of adipose tissue

After transcardial perfusion, adipose tissue was carefully dissected, weighed, and placed in cold PBS on ice. Tissue dissociation was performed using the Adipose Tissue Dissociation Kit (Milenyi Biotec) with minor protocol adjustments. For tissue samples weighing less than 0.5 g, 2.5 mL of dissociation buffer (100 μL of Enzyme D, 25 μL of Enzyme R, and 12.5 μL of Enzyme A in 2.5 mL serum-free DMEM) was used; for larger samples, the volume was adjusted accordingly. Each sample was manually minced into small pieces (2–4 mm) for 60 seconds. The minced tissues were then subjected to two cycles of 20-minute incubation at 37 °C with continuous shaking at 150 rpm followed by running the gentleMACS program “mr_adipose_01”. The resulting cell suspension was applied to a 70 µm cell strainer into a 50 mL tube and washed with 10 mL of DMEM supplemented with 10% FBS. The filtered cells were pelleted by centrifugation at 350 g for 10 minutes and washed three times with PBS.

Cells were immediately stained in LIVE/DEAD Fixable Blue dead cell stain solution (1:1000 in PBS, Invitrogen) for 15 minutes at room temperature and washed three times with PBS. For extracellular staining, cells were incubated on ice in blocking solution (1% Fc block, 5% normal rat serum, and 10% cellBlox in FACS buffer) for 15 minutes, stained with surface antibodies for 30 minutes on ice, and washed three times with FACS buffer. The stained cell pellet was resuspended in 100 µL of Fixation/Permeabilization solution (Invitrogen) and fix overnight at 4 °C. On the following day, fixed cells were washed three times with 1X Permeabilization Buffer (Invitrogen). Cells were then incubated on ice in blocking solution (1% Fc block and 5% normal rat serum in FACS buffer) for 15 minutes, stained with intracellular antibodies for 30 minutes on ice, and washed three times with FACS buffer. Information about antibodies used is provided supplementary tables. Flow cytometry analysis was performed on Cytek Aurora (Cytek Biosciences) and results were analyzed using OMIQ.

### Tissue clearing and immunostaining

Tissue clearing was performed following the AdipoClear protocol^76^. Tissues were fixed in 4% paraformaldehyde (PFA) in PBS overnight at 4 °C and washed three times for 1 hour each.

Following fixation and washes, tissues were dehydrated in graded series of 0%, 20%, 40%, 60%, 80%, and 100% (v/v) methanol/B1n buffer (0.1% Triton-X100, 0.3 M glycine, 0.001% 10N NaOH and 0.01% NaN_3_ in water) with each step lasting 1 hour. Tissues were then delipidated with 100% dichloromethane (DCM) three times (for 1 hour, overnight, and 2 hours, respectively). After two washes with 100% methanol for 1 hour, samples were bleached overnight with 5%

H_2_O_2_ in methanol at 4 °C. Samples were then rehydrated in a reversed methanol/B1n buffer series of 80%, 60%, 40%, 20%, 0% (v/v) methanol/B1n buffer with each step lasting 1 hour. Rehydrated samples were permeabilized in PTxwH Permeabilization Buffer (1X PBS, 0.1% Triton-X100, 0.05% Tween-20, 2 μg/mL heparin, 0.3M glycine, 5% DMSO, and 0.01% NaN_3_) with two 1-hour washes. This was followed by three washes with PTxwH Buffer (1X PBS, 0.1% Triton-X100, 0.05% Tween-20, 2 μg/mL heparin, 0.01% NaN_3_). Permeabilized samples were blocked for 1 day in blocking solution (5% normal goat serum in PTxwH) and then labeled with primary antibodies diluted in blocking solution for 5-7 days in 4°C (anti-RFP, 1:1000, Rockland; anti-MHC class II, 1:200, Abcam). After primary antibody incubation, samples were washed with PTxwH at room temperature for 5 minutes, 10 minutes, 15 minutes, 20 minutes, 1 hour, 2 hours, 4 hours and overnight. Following these washes, samples were incubated in secondary antibodies or dye diluted in blocking solution for 5-7 days in 4°C (anti-rabbit IgG Alexa Fluor 568, 1: 500, Abcam; anti-mouse IgG Alexa Fluor 488, 1: 500, Thermo Fisher Scientific; Hydrazide iFluor 790, 1:500, ATT Bioquest; Isolectin GS-IB4 Alexa Fluor 647, 5 μg/mL, Thermo Fisher Scientific).

After washes, stained tissues were embedded in 1% low melting point agarose (Invitrogen) and dehydrated through a graded series of 25%, 50%, 75%, and three 100% (v/v) methanol/ H_2_O, with each step lasting 1 hour. The samples were then incubated in 100% DCM for three times, each lasting 1 hour. Refractive index matching was achieved by transferring samples to Ethyl Cinnamate solution. Images were acquired using an UltraMicroscope Blaze (Milenyi Biotec).

### Cell-cell interaction analysis

To identify cell-cell communication events that changed linearly with age, we first generated pseudo-bulk profiles and filtered samples with fewer than 10 cells or 1000 counts. After filtering, we used PyDESeq2 to perform differential expression analysis (DEA) for each cell type with age as a continuous independent variable. Interaction statistics were calculated from the DEA results using the “li.multi.df_to_lr” function from the Liana package ^77^. We utilized consensus ligand-receptor interaction resources, and interactions expressed in less than 10% of the nuclei per cell type in 32-month-old samples were filtered out. Ligand-receptor interactions with an adjusted *p-value* of less than 0.05 were considered to have significantly changed with age.

### Linear modeling

We performed differential expression analysis using pyDESeq2, fitting regressions to extract the individual effects of age, adipocyte number, and adipocyte size on the pseudo-bulk gene expression of each cell type. The model used was Gene ∼ age + adipocyte number + adipocyte size. For each variable, GSEA was performed on the Wald statistics obtained from the DESeq2 results to identify significantly enriched MSigDB Hallmark gene sets associated with that variable. To validate our results, we calculated activity of “IFNγ response” and “Inflammation response” using Scanpy’s “sc.tl.score_genes function”. The average score across all adipocytes was calculated for each sample. Subsequently, we employed linear regression to model the relationship between these scores with our independent variables. To visually assess influence of individual predictors, we generated component-plus-residual plots (also known as C+R plots or partial residual plots) using crPlots().

### Isolation and culturing of primary ASPCs

Primary mouse ASPCs were isolated from subcutaneous adipose tissues of young mice. Briefly, dissected adipose tissue was manually minced into small pieces (1–3 mm) and digested in 5 mL of 1 mg/mL (0.1%) collagenase II solution in DMEM for 60 minutes at 37 °C with continuous shaking at 150 rpm. Following digestion, 13 mL of complete media (10% FBS and 1X anti-Anti in DMEM) was added to stop the reaction, and the cell suspension was filtered through a 100 µm cell strainer. The filtered cells were pelleted by centrifugation at 350 g for 10 minutes and washed twice by resuspending in complete media followed by centrifugation. After washes, cells from each animal were resuspended in 2 mL of complete media and seeded in one well of a 6-well collagen-coated plate. Primary ASPCs were cultured in a humidified 37 °C incubator with 5% O_2_ and 5% CO_2_. On the following day, adherent preadipocytes were washed, trypsinized, and replated onto a 10 cm collagen-coated plate to reduce potential contamination of other cell types in the stromal vascular fraction. All experiments involving ASPCs were performed using cells between passage 3 and 4.

### Characterization of the senescence phenotype in cultured ASPCs

We characterized the development of a senescence phenotype in primary ASPCs across four independent experiments. Primary ASPCs were collected from four different animals, cultured for a minimum of ten days, exposed to 10-Gy X-rays, split two days post-irradiation, and supplied with fresh media every two to three days. Irradiated ASPCs were collected at various time points after irradiation to measure the expression of cellular senescence markers, while non-irradiated ASPCs served as controls.

For RNA sequencing experiments, RNA was extracted using the RNeasy Plus Mini Kit (Qiagen). Sequencing libraries were prepared using the NEBNext Ultra II Directional RNA Library Prep Kit (New England Biolabs). Sequencing was performed using a NovaSeq 6000 Sequencing system (Illumina) with 150 paired-end reads. Raw sequencing reads were demultiplexed into FASTQ format files using bcl2fastq (Illumina) and aligned to the GRCm39 mouse reference genome using STAR (version 2.7.10)^78^. Gene counts were produced using HTseq (version 2.0.5) with default settings except “-m intersection-strict”^79^. Normalization and differential expression analysis were performed using DESeq2. GSEA was performed on the DESeq2 results using GSEApy^80^. To examine the expression changes of secreted proteins, we curated genes annotated to be in the “extracellular space” with a confidence greater than 3 stars in the COMPARTMENTS database^81^.

Immunocytochemistry was performed using standard protocols. Briefly, ASPCs were seeded onto 8-well chamber slides (Nunc™ Lab-Tek™ II Chambered Coverglass) and fixed at various time points after irradiation. Fixation was performed by incubating PBS-washed cells in 4% PFA in PBS at room temperature for 15 minutes. Fixed cells were washed with PBS, permeabilized with 0.5% Triton X-100 in PBS for 10 minutes and blocked with blocking buffer (5% normal goat serum in PBS with 0.1% Triton) for 1 hour at room temperature. Cells were then incubated with primary antibodies (anti-p16^INK4a^, 1:100, Abcam; anti-LaminB1, 1:500, Santa Cruz) diluted in blocking buffer for 1 hour at room temperature. Cells were washed three times with wash buffer (PBS with 0.1% Triton) and incubated with secondary antibodies (anti-rabbit IgG Alexa Fluor™ 488, 1:250, Invitrogen; goat-anti-mouse IgG Alexa Fluor™ 488, 1:250, Invitrogen) diluted in blocking buffer for 1 hour in the dark at room temperature. Cells were washed, stained in DAPI solution (1 mg/mL in PBS) for 14 minutes in the dark at room temperature, washed again, and stored in PBS prior to imaging. For cell proliferation measurements, cells were labeled with 10 μM of EdU diluted in fresh complete media for 24 hours prior to fixation. EdU was detected using the Click-iT™ Plus EdU Cell Proliferation Kit for Imaging (Invitrogen) following manufacturer’s protocol. Microscopy images were acquired with a Nikon Spinning Disk confocal microscope. Fluorescence intensity was measured from segmented nuclei using ImageJ.

### Senescent cell transplantation

3-month-old male C57BL/6J mice, purchased from the Jackson Laboratory, were acclimated for one-week, and baseline measurements were collected prior to senescent cell transplantation.

Mice were randomly assigned to either the control or senescent cell recipient group by a researcher not involved in data collection. The randomization was stratified to ensure no significant differences in baseline measurements (including rotarod performance, body weight, body composition, and glucose tolerance) between the final experimental groups. On the day of transplantation, senescent or control ASPCs were collected by trypsinization. Cells were washed twice by resuspending in PBS followed by centrifugation at 350 g for 5 minutes. Washed cells were resuspended in PBS at 0.5×10^7^ cells/mL. Each mouse was injected intraperitoneally (IP) with 1×10^6^ control or senescent cells in 200 μL of PBS using 26-gauge needles (we have confirmed that passing cells through 26-gauge needles did not interfere with the viability of control and senescent ASPCs). The investigator responsible for all subsequent behavioral and physiological measurements was blinded to the experimental group assignments. Blinding was maintained throughout the study until the day of tissue collection. Visceral adipose tissues were collected at one-month post-transplantation for flow cytometry analysis and at two-months post-transplantation for single-nucleus RNA sequencing.

### Longitudinal measurement and localization of transplanted ASPCs

For longitudinal tracking of transplanted ASPCs, mice were injected with 1×10^6^ senescent or control ASPCs derived from young *Actin-Cre; Rosa26LSL-OVA-Luc* mice. Bioluminescence imaging (BLI) was performed using the Lago imaging system (Spectral Instruments Imaging) at two, five, seven, fourteen, twenty-one- and twenty-eight-days post-transplantation. Mice were anesthetized and injected intraperitoneally with warm D-luciferin solution (150 mg/kg). From 8 to 16 minutes post-D-luciferin injection, an image was captured every minute with 50-second exposure and medium binning. Maximum background-corrected bioluminescence of each mouse was measured using Aura Image software (Spectral Instruments).

To identify the location of transplanted ASPCs, mice were injected with 1×10^6^ senescent or control ASPCs derived from young *CAG-mRFP1/J* mice. At two, five, and seven days post-injection, recipient mice and a non-injected negative control mouse were euthanized. Organs within the abdominal cavity, including the peritoneum, kidney, spleen, liver, testis, pancreas, the digestive tract (from stomach to distal colon), and various adipose tissues (perigonadal, mesenteric, and perirenal), were collected. Each organ was examined carefully under a fluorescence dissection microscope. We consistently observed RFP signal in perigonadal adipose tissues within the first week after transplantation. To validate the presence of transplanted cells, tissue clearing and immunostaining were performed on perigonadal adipose tissues from recipient mice (see “Tissue clearing and immunostaining” for details).

### Rotarod assay

Rotarod performance was measured using an accelerating RotaRod system (Ugo Basile) following a previously described protocol^82^. Mice were initially trained on the rotarod at constant speeds of 4, 6, 8 rpm, with each session lasting 300 seconds and separated by at least a 30-minute break. Rotarod performance was then measured before ASPC transplantation, and at 1-and 2-months post-transplantation. On measurement days, mice were acclimatized to the room for at least 30 minutes. The rotating speed was set to accelerate from 4 to 40 rpm over 300 seconds. The time to fall for each mouse was recorded, and the average across three trials was used for each mouse.

### Body composition measurement

Body composition of recipient mice was measured using an EchoMRI machine before ASPC transplantation, and at 1- and 2-months post-transplantation. Mice were weighed and body fat mass and lean mass were determined following the manufacturer’s protocol.

### Glucose tolerance test

A glucose tolerance test (GTT) was performed on recipient mice before ASPC transplantation and at 1-month post-transplantation. Mice were placed in a clean cage without food and with *ad libitum* access to water starting at 5:00 pm the night prior to the test. On the following morning at 9:00 am, the weight and fasting blood glucose level of each mouse were measured. Each mouse was then injected intraperitoneally with 20% glucose solution at a dose of 1.5 g/kg. Blood glucose levels were subsequently measured at 15, 30, 60, and 120 minutes after glucose injection.

### MHC class II induction in primary endothelial cells

Primary mouse aortic, dermal lymphatic, and liver sinusoidal endothelial cells were purchased from Cell Biologics Inc. Cells were plated on collagen-coated culture dishes and expanded in complete mouse endothelial cell medium (Cell Biologics, catalog no. M1168). Fresh media with recombinant mouse IFNγ (50 ng/mL) or 48-hour conditioned media from control or senescent ASPCs were added to the endothelial cells. After 24, 48, or 72 hours of treatment, or after 72 hours of treatment followed by a 72-hour washout period, cells were collected by trypsinization, stained, and analyzed by flow cytometry.

### CD4^+^ T cell co-culture with endothelial cells

Naïve CD4^+^ T cells were isolated from spleens of 4-month-old OT-II mice using the EasySep™ Mouse Naïve CD4^+^ T Cell Isolation Kit (Stemcell Technologies) following manufacturer’s instructions. Isolated naïve CD4^+^ OT-II cells were stained with Cell Proliferation Dye eFluor™ 670 (Invitrogen) at 1× 10^6^ cells/mL in 5 µM dye diluted in PBS. Primary mouse endothelial cells were pre-treated for 72 hours with the cognate antigen peptide OVA_323-339_ (1 µg/mL) and recombinant mouse IFNγ (50 ng/mL). Immortalized Mouse Dendritic Cells (MutuDC1940) pre-treated with OVA_323-339_ (1 µg/mL) were used as a positive control.

1× 10^5^ stained naïve CD4^+^ OT-II cells were added to each well of a 96-well plate containing pre-treated endothelial cells or dendritic cells. After 72 hours of co-culture, T cells were collected, stained, and analyzed by flow cytometry.

### Vessel-on-a-chip experiment

The vessel-on-a-chip microfluidic system was manufactured based on a previous protocol^83^. Microfluidic channel was designed using SolidWorks (1 mm width and height, 5 mm length, and inlet/outlet reservoir diameters of 1.5 mm and 2 mm, respectively). The master mold was fabricated using photolithography by coating a silicon wafer with 1 mm thick SUEX dry film photoresist (iMicro Materials, Inc.). Polydimethylsiloxane (PDMS, Dow Corning) base and cross-linker were mixed at a 10:1 ratio, poured onto the mold, and cured at 80 for 2 hours.

Inlets and outlets of the PDMS device were punched with a 1 mm biopsy punch. The devices were plasma bonded to PDMS-coated glass slides and incubated at 80 for 30 minutes to enhance binding. Devices were pretreated by filling with 10% v/v (3-aminopropyl)triethoxysilane (Sigma-Aldrich) in ethanol for 15 minutes at room temperature. They were washed extensively with ethanol and dried at 80 for 2 hours. Gravitational lumen patterning (GLP) was used to create the 3D hydrogel lumens. Briefly, after degassing the devices in a vacuum chamber, they were filled with an ice-cold hydrogel mixture consisting of 110 µL of rat tail type I collagen (10 mg/mL, Corning), 40 μL HEPES (1 M, Gibco), 14 µL sodium bicarbonate (1 M, ThermoFisher Scientific), 3 µL sodium hydroxide (1 M, ThermoFisher Scientific) and 33 µL of Complete Endothelial Cell Medium (Cell Biologics). Devices were then rotated and perfused with 50 µL of ice-cold cell media to form 3D lumens within the hydrogel and allowed to fully polymerize for one day in an incubator.

Primary mouse aortic endothelial cells (6 × 10^6^ cells/mL) were seeded inside the devices in four sequential steps, each with a 50-minute incubation period. This allowed for cell adherence to the four sides of the developing lumen. The devices were kept in a humidified 37 °C incubator with 5% CO2 and supplied with fresh media daily throughout the experimental duration. After confluent endothelial vessels were established (typically two days post-seeding), media containing either OVA_323-339_ (1 µg/mL), recombinant mouse IFNγ (50 ng/mL), or both OVA_323-339_ and IFNγ were added for 72 hours to induce MHC II expression. CD4^+^ T cells were isolated from spleens of 4-month-old OT-II mice using EasySep™ Mouse CD4^+^ T Cell Isolation Kit (Stemcell Technologies) and stained with Cell Proliferation Dye eFluor™ 670 (Invitrogen).

Before the addition of T cells, vessels were thoroughly washed with fresh media to remove residual antigen and IFNγ. 1 × 10^5^ stained T cells were added to the inlet of each device and allowed to interact with the vessel for 24 hours. After 24 hours, z-stack images of the devices were acquired using a Nikon Spinning Disk confocal microscope to determine the total number of T cells inside each vessel. The vessels were then thoroughly washed with PBS to remove T cells sitting on top of the endothelial cells. Washed vessels were subsequently fixed with 4% PFA in PBS for 15 minutes, washed three times with PBS, permeabilized with 0.01% Tween-20 in PBS, and stained with WGA-Alexa Fluor 488 (1:200) and DAPI (1:2000) for 1.5 hours. Following extensive washes, z-stack images of the stained devices were taken using a Nikon Spinning Disk confocal microscope.

Image analysis was performed in ImageJ. To count T cells, thresholding using the “Triangle” method was performed on the Z projection of the far-red channel. Binary dilation and watershed operations were then applied. The “analyze particles” function was used to identify and count cells. The percentage of cells retained by or migrated through the vessel were calculated by dividing the number of cells after staining by the number of cells before washes in the same field of view.

### T cell depletion from aged mice

20-month-old female or 21-month-old male C57BL/6J mice, purchased from the Jackson Laboratory, were acclimated for one-week prior to antibody treatment. Mice were randomly assigned to either the IgG2b control (BioXCell, #BE00090), anti-mouse CD4 (BioXCell, #BE0003-1), or anti-mouse CD8β (BioXCell, #BE0223) group. Each mouse received an intraperitoneal injection 5 mg/kg antibody twice per week for a total of ten doses. T cell depletion in peripheral blood was confirmed a week prior to tissue collection. Only the IgG2b control and anti-mouse CD4 were used for single-nuclei RNA sequencing.

### Reporting summary

Further information on research design is available in the Nature Portfolio Reporting Summary linked to this article.

---

**Extended Data Fig. 1.**
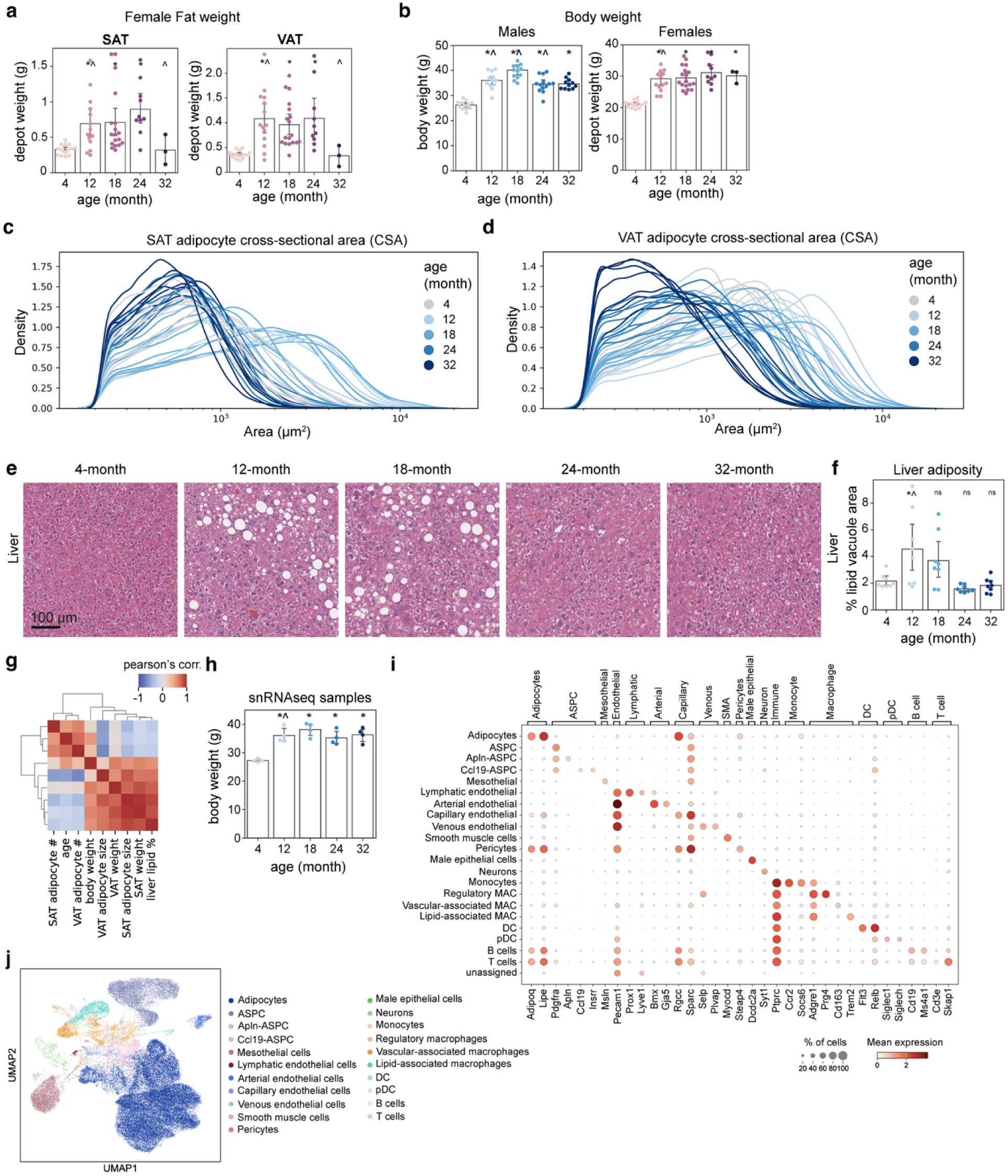
| Histological characterization of tissue remodeling and cell type annotation. **a**,**b**, Weight of SAT and VAT in female mice (**b**) and body weight for both sexes (**b)** across five age groups (*n* = 3–18 per group). **c**,**d**, Frequency distribution of adipocyte cross-sectional area (CSA) in SAT (**c**) and VAT (**d**). Each trace represents the distribution from an individual animal, colored by age (*n* = 7–8 per group). **e**,**f**, Representative H&E-stained liver sections (**e**) and corresponding quantification of lipid vacuole area (**f**) across five age groups (*n* = 8 per group). **g**, Heatmap of Pearson’s correlation coefficients between age, body weight, adipose tissue weight, adipocyte number, adipocyte size and liver lipid area, showing a positive correlation between liver lipid deposition and adiposity. **h**, Body weight of the subset of mice used for single-nucleus RNA sequencing (*n* = 4 per group). **i**, Dot plot showing the expression of canonical marker genes used for cell type annotation. **j**, UMAP projections of sequenced nuclei from SAT and VAT, colored by further refined cell subtypes. In all bar plots (a, b, f and h), data are presented as mean ± 95% confidence intervals, with each dot representing an individual mouse. Statistical significance was determined by one-way ANOVA with Tukey’s post hoc test or Welch’s two-sided t-test when variances were not equal. Asterisks (*) indicate p-value < 0.05 compared to 4-month-old mice, and carets (^) indicate p-value < 0.05 compared to the preceding age group.

**Extended Data Fig. 2.**
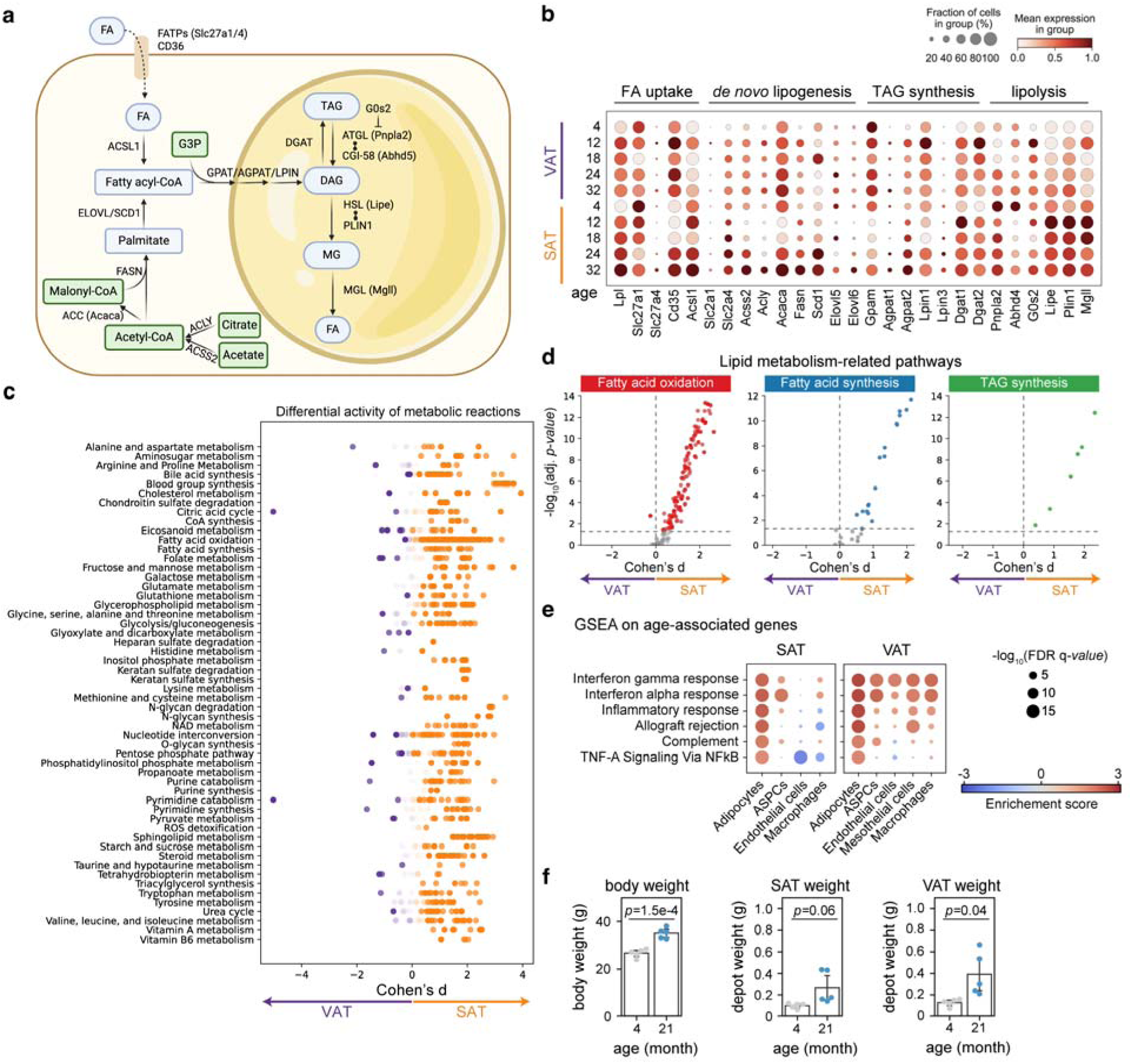
| Depot-specific and age-dependent metabolic and inflammatory remodeling in adipocytes. **a**, Schematic showing the major lipid metabolism pathways in the adipocyte. **b**, Dot plot showing the expression of individual genes involved in fatty acid uptake, *de novo* lipogenesis, triacylglyceride (TAG) synthesis and lipolysis in adipocytes across all age groups. **c**, COMPASS analysis of differential metabolic reaction activity between SAT and VAT adipocytes from 4-month-old mice. Reactions are grouped by metabolic subsystems. Orange dots indicate higher predicted metabolic activity in SAT, and purple dots indicate higher activity in VAT. **d,** Predicted activity of lipid metabolism reactions in adipocytes. **e,** Dot plot summarizing the age-related enrichment of inflammation-related gene sets across major cell types from SAT and VAT. Scores were adjusted for tissue weight to isolate age-associated signals. A positive score (red) indicates increased expression with age. **f,** Body and adipose tissue weight of 4- and 21-month-old mice used for flow cytometry.

**Extended Data Fig. 3.**
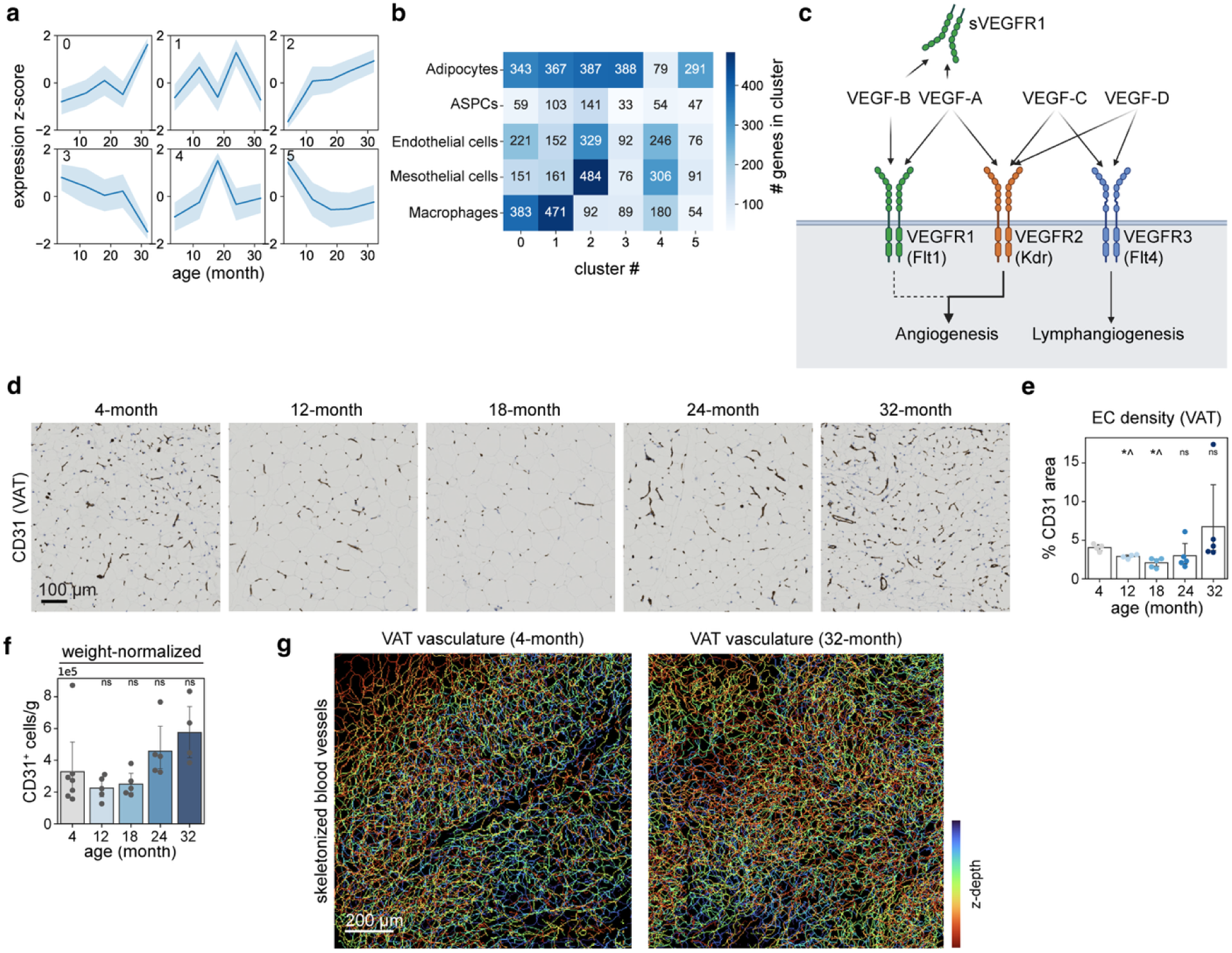
| Temporal gene expression trajectories and age-related vascular remodeling. **a**,**b**, Clustering of genes into six major temporal expression trajectories in VAT (**a**) and the relative contribution of individual cell types to each gene cluster (**b**). **c**, Schematic showing the VEGF signaling pathways involved in blood and lymphatic vessel proliferation. **d**,**e**, Representative images (**d**) and quantification (**e**) of vascular density in VAT across all age groups (*n* = 4–5 per group). **f**, Flow cytometric quantification of blood endothelial cell density in VAT, normalized to tissue weight (*n* = 4–7 per group). **g**, Representative images of skeletonized vasculature derived from Isolectin-IB4 staining of whole-mount VAT from 4- and 32-month-old mice. While total endothelial cell numbers decline in late life, the slower rate of decline relative to lipid loss suggests that increased vascular density likely results from the slow pruning of existing vessels rather than active angiogenesis. In all bar plots (e and f), data are presented as mean ± 95% confidence intervals, with each dot representing an individual mouse. Statistical significance was determined by Welch’s two-sided t-test. Asterisks (*) indicate p-value < 0.05 compared to 4-month-old mice, and carets (^) indicate p-value < 0.05 compared to the preceding age group.

**Extended Data Fig. 4.**
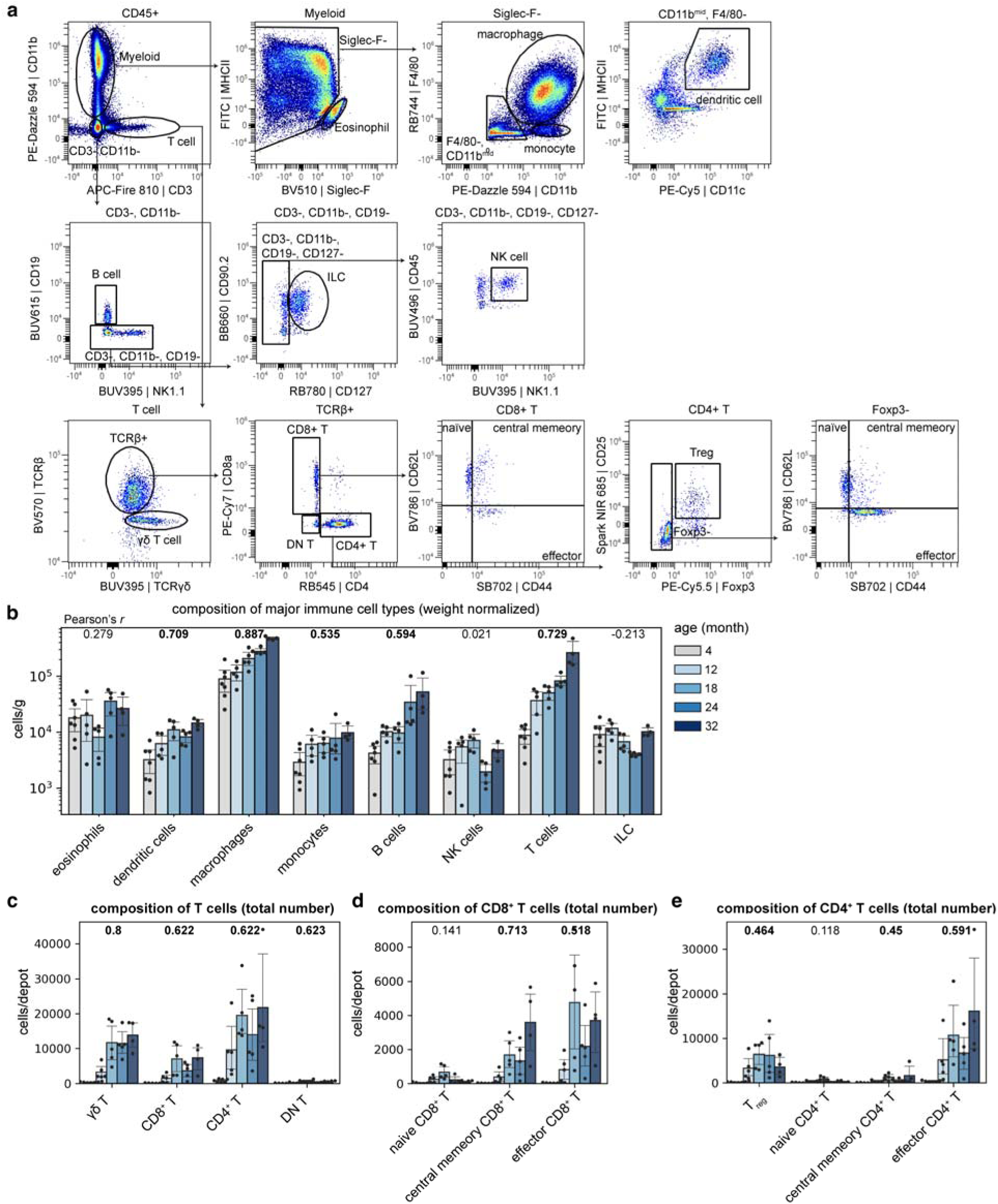
| Age-related immune cell composition changes in VAT. **a**, Flow cytometry gating strategies for the identification and quantification of major immune cell populations in adipose tissue. **b**, Tissue weight-normalized density of major immune cell types in VAT across all five age groups. **c–e**, Quantification of the total number of T-cell subtypes, including all T cells (**c**), CD8^+^ T cells (**d**) and CD4^+^ T cells (**e**). In all bar plots (b–e), data are presented as mean ± 95% confidence intervals, with each dot representing an individual mouse (*n* = 4–7 per group). Pearson’s correlation coefficients (ρ) with age are shown; bolded values indicate a significant correlation (*p*-value < 0.05).

**Extended Data Fig. 5.**
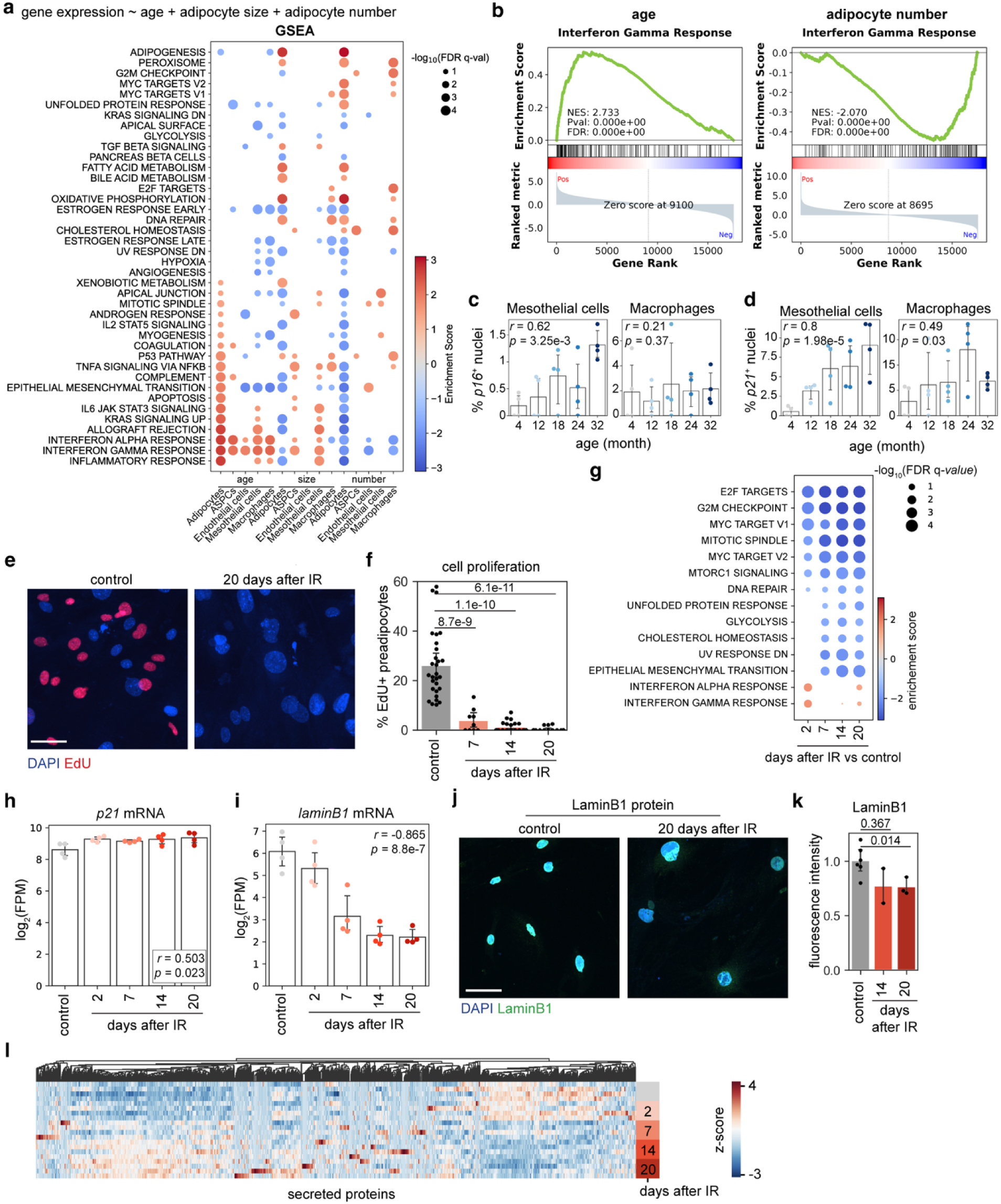
| Dissecting the age-related IFNγ response and characterization of irradiation-induced senescent ASPC. **a**, Dot plots summarizing gene sets significantly associated with age, adipocyte size and adipocyte number across major cell types in VAT. The independent effect of each variable after correcting for the other two is shown. A positive score (red) indicates a positive correlation. Dot size corresponds to the FDR *q*-value. **b**, GSEA enrichment plots showing the association of IFNγ response genes with age (left) or adipocyte number (right) in adipocytes. **c**,**d**, Percentage of *p16*-expressing (**c**) and *p21*-expressing (**d**) nuclei in various cell types in VAT and SAT across all age groups. Each dot represents a mouse (*n* = 4 per group). **e**,**f**, Representative images (**e**) and quantification (**f**) of ASPC proliferation based on EdU incorporation at specified timepoints following irradiation. **g**, Dot plot summarizing gene sets enriched in ASPCs at different days after irradiation compared to unirradiated controls. **h**,**i**, Expression of *p21* and *laminB1* over time in ASPCs following irradiation (*n* = 4 per timepoint). **j**,**k**, representative image (**j**) and quantification (**k**) of laminB1 protein levels in control versus senescent ASPCs (*n* = 3–5 per timepoint). **l**, Heatmap showing transcriptional changes in secreted proteins, suggesting the development of a robust Senescence-Associated Secretory Phenotype (SASP) in irradiated ASPCs. In all bar plots (**c**, **d**, **f**, **h**, **i** and **k**), data are presented as mean ± 95% confidence intervals, with each dot representing an independent biological replicate (*n* = 3–6 per timepoint). Statistical significance was determined by Welch’s two-sided t-test or Pearson’s correlation.

**Extended Data Fig. 6.**
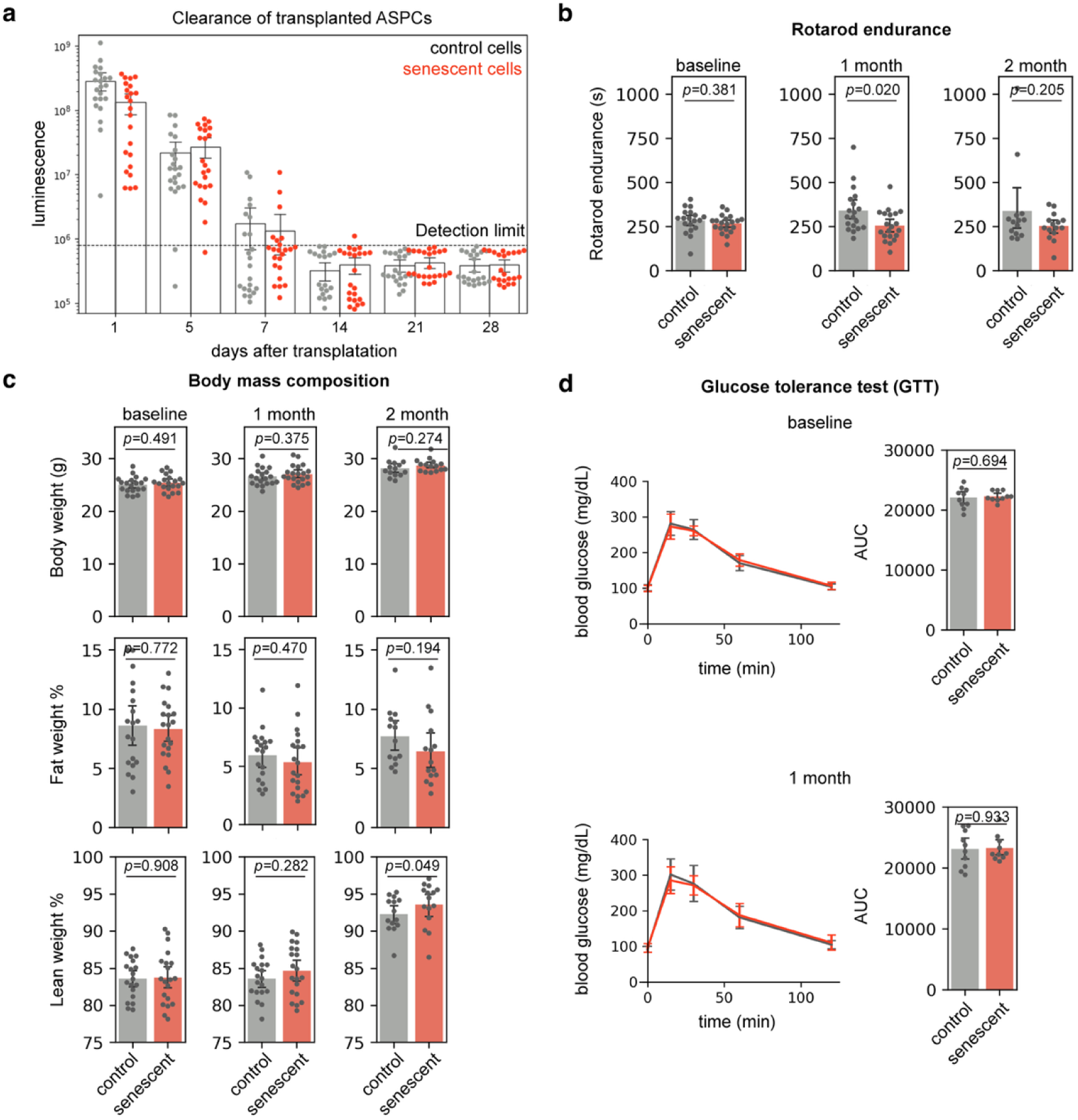
| Physiological and functional assessment of senescent-cell recipient mice. **a**, Longitudinal *in vivo* bioluminescence imaging tracking the clearance of luciferase-expressing transplanted ASPCs (*n* = 22–23 per group). **b,c**, Rotarod performance (**b**) and body mass composition (**c**) of recipient mice measured at baseline and at one and two months after transplantation (*n* = 14–20 per group). **d**, Glucose tolerance test performed on recipient mice measured at baseline and one month after transplantation (*n* = 10 per group). In all plots, data are presented as mean ± 95% confidence intervals, with each dot represents an individual mouse. Statistical significance was determined by Welch’s two-sided t-test.

**Extended Data Fig. 7.**
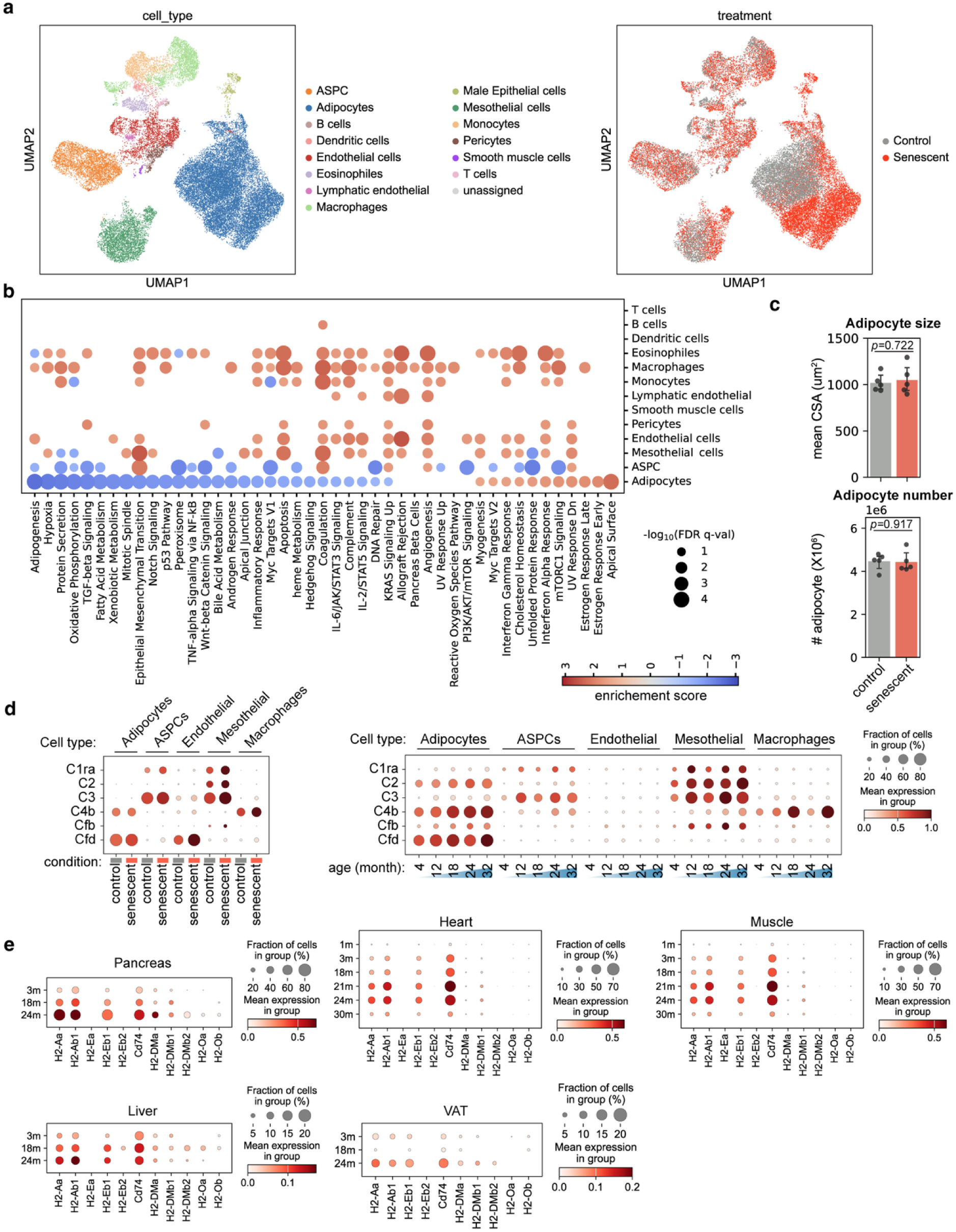
| Transcriptomic changes in VAT after senescent cell transplantation. **a**, UMAP projection of sequenced nuclei from the VAT of young recipient mice at two months post injection, colored by cell type (left) and treatment group (right). **b**, GSEA results showing the effect of senescent cell exposure across various VAT cell types. A positive enrichment score (red) indicates higher expression in the senescent cell recipient group. **c**, No significant change in mean adipocyte CSA (top) and total adipocyte number (bottom) in VAT at two months after senescent cell injection. **d**, Expressions of complement system components across different cell types in VAT after control or senescent cell transplantation or at different ages. **e**, Expression of MHC II molecules in endothelial cells across various organs (pancreas, heart, skeletal muscle, liver, VAT) from the Tabula Muris Senis dataset.

**Extended Data Fig. 8.**
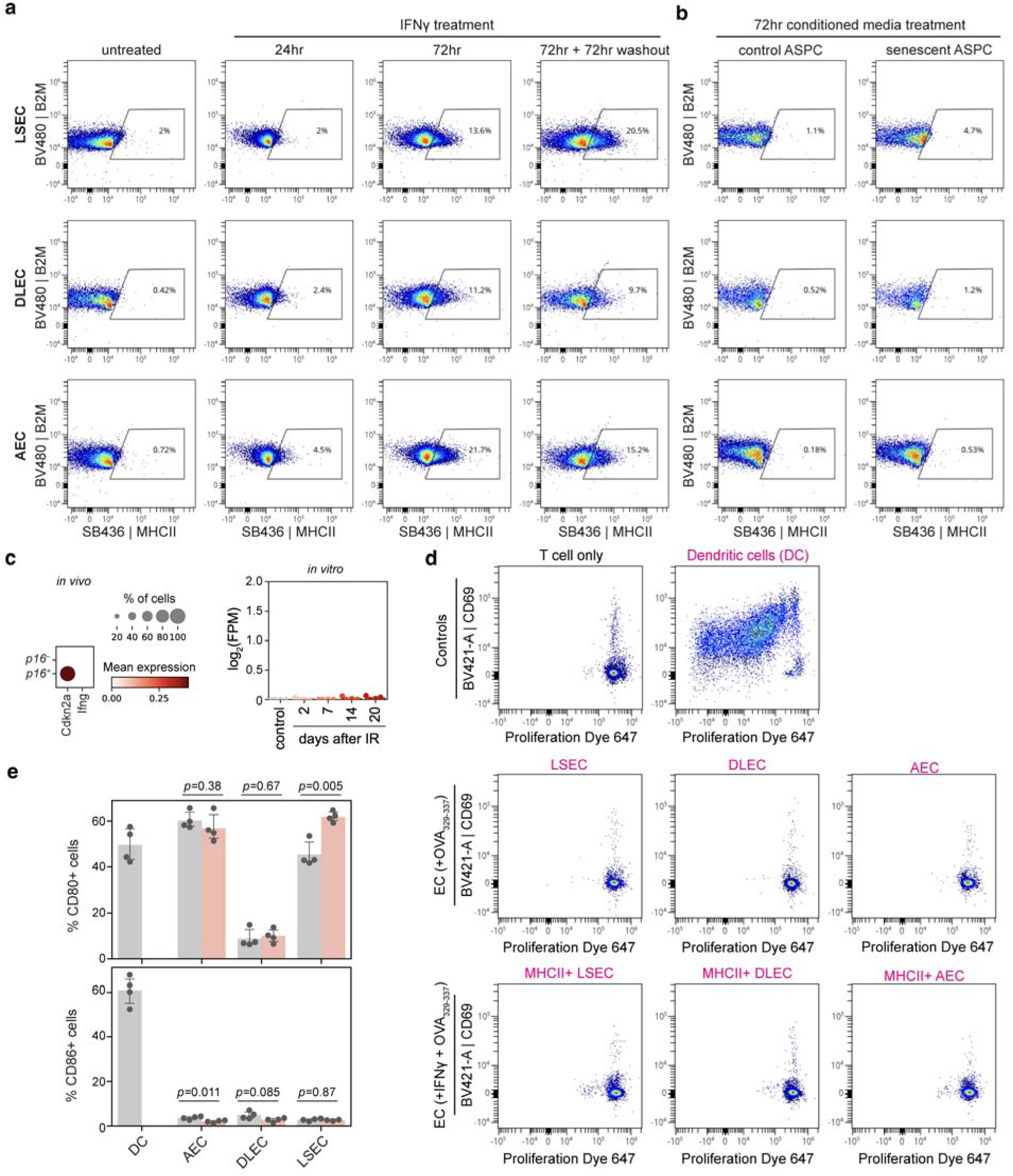
| Characterization of endothelial MHC II induction and function. **a**,**b**, Representative flow cytometry plots showing MHC class I and class II expression on primary endothelial cells following treatment with recombinant IFNγ (**a**) or with conditioned media from control versus senescent ASPCs (**b**). **c**, expression of *Ifng* in senescent cells identified *in vivo* (adipose tissue) and *in vitro* (irradiated ASPCs). **d**, Representative flow cytometry plots showing the activation and proliferation status of naïve CD4^+^ OT-II cells after 72 hours of co-culture with either professional antigen-presenting cells (dendritic cells) or MHC II-expressing endothelial cells. **e**, Quantification of the percentage of cells expressing the co-stimulatory molecules CD80 and CD86 with or without IFNγ treatment. Data are presented as mean ± 95% confidence intervals, with each dot representing an independent biological replicate (*n* = 4 per group). Statistical significance was determined by Welch’s two-sided t-test.

**Extended Data Fig. 9.**
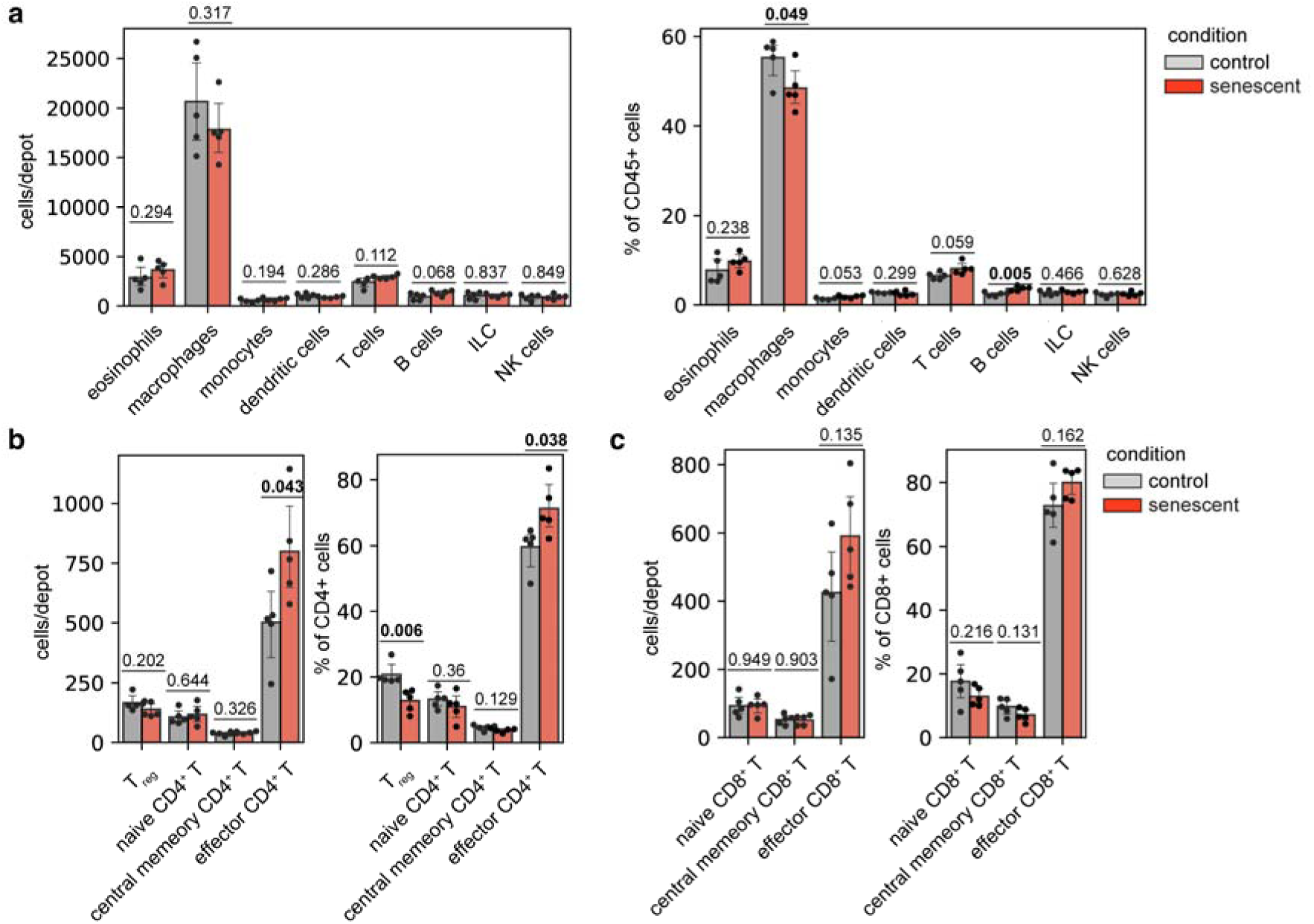
| Immune cell composition changes in young VAT following senescent cell transplantation. **a–c**, Absolute cell number (left) and relative proportions (right) of major immune cell populations (**a**), CD4^+^ T cell subsets (**b**) and CD8^+^ T cell subsets (**c**) in VAT of young recipient mice. Data are presented as mean ± 95% confidence intervals, with each dot representing an individual mouse (*n* = 5 per group). Statistical significance was determined by Welch’s two-sized t-test.

**Extended Data Fig. 10.**
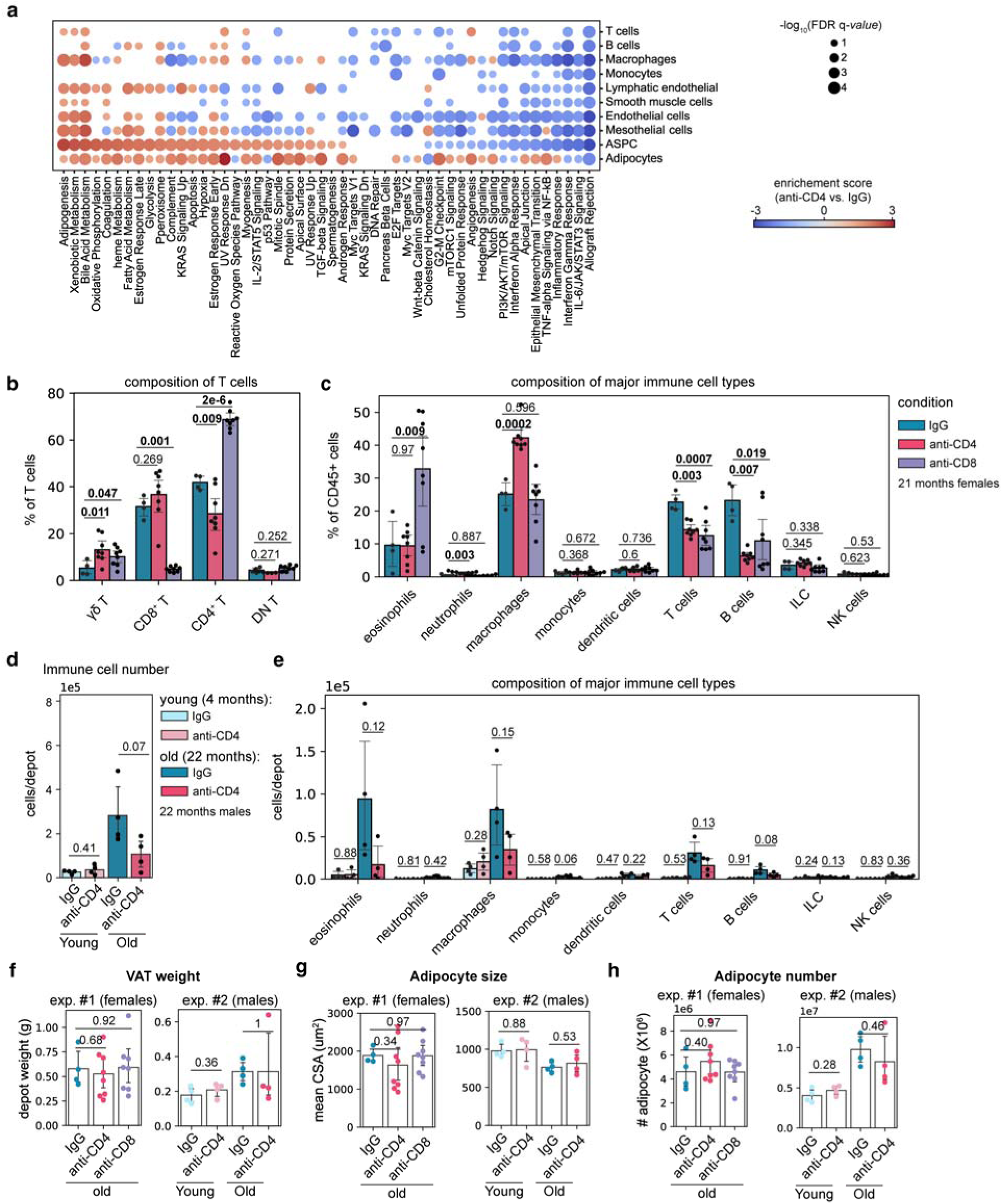
| Impact of T-cell depletion on young and aged VAT. **a**, GSEA results showing the transcriptomic impact of CD4^+^ T-cell depletion across various cell types in aged VAT compared to isotype IgG controls. **b**,**c**, Relative proportions of T cell subtypes (**b**) and major immune cell types (**c**) in VAT of aged female mice after CD4^+^ or CD8^+^ depletion. **d**,**e**, Absolute numbers of total immune cells (**d**) and major immune subtypes (**e**) in VAT of young or aged male mice after CD4^+^ T-cell depletion. **f**–**h**, No significant changes in tissue weight (**f**), mean adipocyte CSA (**g**) or total adipocyte number (**h**) in VAT after T-cell depletion. For all bar plots (**b**–**h**), data are presented as mean ± 95% confidence intervals, with each dot representing an individual mouse (*n* = 4–8 per group). Statistical significance was determined by Welch’s two-sized t-test.

## References

1 Ou, M. Y., Zhang, H., Tan, P. C., Zhou, S. B. & Li, Q. F. Adipose tissue aging: mechanisms and therapeutic implications. Cell Death Dis 13, 300 (2022). 10.1038/s41419-022-04752-6

2 Nguyen, T. T. & Corvera, S. Adipose tissue as a linchpin of organismal ageing. Nat Metab 6, 793–807 (2024). 10.1038/s42255-024-01046-3

3 Liu, J., Huang, Q. & Liu, F. Fat talks first: how adipose tissue sets the pace of aging? Life Med 4, lnaf028 (2025). 10.1093/lifemedi/lnaf028

4 Rosen, E. D. & Spiegelman, B. M. What we talk about when we talk about fat. Cell 156, 20–44 (2014). 10.1016/j.cell.2013.12.012

5 Chouchani, E. T. & Kajimura, S. Metabolic adaptation and maladaptation in adipose tissue. Nat Metab 1, 189–200 (2019). 10.1038/s42255-018-0021-8

6 Trim, W. V. & Lynch, L. Immune and non-immune functions of adipose tissue leukocytes. Nat Rev Immunol 22, 371–386 (2022). 10.1038/s41577-021-00635-7

7 Kahn, C. R., Wang, G. & Lee, K. Y. Altered adipose tissue and adipocyte function in the pathogenesis of metabolic syndrome. J Clin Invest 129, 3990–4000 (2019). 10.1172/JCI129187

8 Frasca, D., Blomberg, B. B. & Paganelli, R. Aging, Obesity, and Inflammatory Age-Related Diseases. Front Immunol 8, 1745 (2017). 10.3389/fimmu.2017.01745

9 Hotamisligil, G. S. Inflammation and metabolic disorders. *Nature* **444**, 860–867 (2006). 10.1038/nature05485

10 Stout, M. B., Justice, J. N., Nicklas, B. J. & Kirkland, J. L. Physiological Aging: Links Among Adipose Tissue Dysfunction, Diabetes, and Frailty. Physiology (Bethesda) 32, 9–19 (2017). 10.1152/physiol.00012.2016

11 Neto, A., Fernandes, A. & Barateiro, A. The complex relationship between obesity and neurodegenerative diseases: an updated review. Front Cell Neurosci 17, 1294420 (2023). 10.3389/fncel.2023.1294420

12 Murphy, R. A. et al. Weight change, body composition, and risk of mobility disability and mortality in older adults: a population-based cohort study. J Am Geriatr Soc 62, 1476–1483 (2014). 10.1111/jgs.12954

13 Dorling, J. L., Martin, C. K. & Redman, L. M. Calorie restriction for enhanced longevity: The role of novel dietary strategies in the present obesogenic environment. Ageing Res Rev 64, 101038 (2020). 10.1016/j.arr.2020.101038

14 Soukas, A. A., Hao, H. & Wu, L. Metformin as Anti-Aging Therapy: Is It for Everyone? Trends Endocrinol Metab 30, 745–755 (2019). 10.1016/j.tem.2019.07.015

15 Xu, L. et al. PPARgamma agonists delay age-associated metabolic disease and extend longevity. Aging Cell 19, e13267 (2020). 10.1111/acel.13267

16 Hickson, L. J. et al. Senolytics decrease senescent cells in humans: Preliminary report from a clinical trial of Dasatinib plus Quercetin in individuals with diabetic kidney disease. EBioMedicine 47, 446–456 (2019). 10.1016/j.ebiom.2019.08.069

17 Al Qassab, M. et al. The Expanding Role of GLP-1 Receptor Agonists: Advancing Clinical Outcomes in Metabolic and Mental Health. Curr Issues Mol Biol 47 (2025). 10.3390/cimb47040285

18 Kuk, J. L., Saunders, T. J., Davidson, L. E. & Ross, R. Age-related changes in total and regional fat distribution. Ageing Res Rev 8, 339–348 (2009). 10.1016/j.arr.2009.06.001

19 Petr, M. A. et al. A cross-sectional study of functional and metabolic changes during aging through the lifespan in male mice. Elife 10 (2021). 10.7554/eLife.62952

20 Palliyaguru, D. L. et al. Fasting blood glucose as a predictor of mortality: Lost in translation. Cell Metab 33, 2189–2200 e2183 (2021). 10.1016/j.cmet.2021.08.013

21 Ferrucci, L. & Fabbri, E. Inflammageing: chronic inflammation in ageing, cardiovascular disease, and frailty. Nat Rev Cardiol 15, 505–522 (2018). 10.1038/s41569-018-0064-2

22 Schaum, N. et al. Ageing hallmarks exhibit organ-specific temporal signatures. Nature 583, 596-602 (2020). 10.1038/s41586-020-2499-y

23 Ouchi, N., Parker, J. L., Lugus, J. J. & Walsh, K. Adipokines in inflammation and metabolic disease. Nat Rev Immunol 11, 85–97 (2011). 10.1038/nri2921

24 Kawai, T., Autieri, M. V. & Scalia, R. Adipose tissue inflammation and metabolic dysfunction in obesity. Am J Physiol Cell Physiol 320, C375–C391 (2021). 10.1152/ajpcell.00379.2020

25 Tchkonia, T. et al. Fat tissue, aging, and cellular senescence. Aging Cell 9, 667–684 (2010). 10.1111/j.1474-9726.2010.00608.x

26 Krotkiewski, M., Bjorntorp, P., Sjostrom, L. & Smith, U. Impact of obesity on metabolism in men and women. Importance of regional adipose tissue distribution. J Clin Invest 72, 1150–1162 (1983). 10.1172/JCI111040

27 Wang, Q. A., Tao, C., Gupta, R. K. & Scherer, P. E. Tracking adipogenesis during white adipose tissue development, expansion and regeneration. Nat Med 19, 1338–1344 (2013). 10.1038/nm.3324

28 Tabula Muris, C. A single-cell transcriptomic atlas characterizes ageing tissues in the mouse. Nature 583, 590–595 (2020). 10.1038/s41586-020-2496-1

29 Wu, Y. et al. Dynamics of single-nuclei transcriptomic profiling of adipose tissue from diverse anatomical locations during mouse aging process. Sci Rep 14, 16093 (2024). 10.1038/s41598-024-66918-w

30 Zhang, Z. et al. A panoramic view of cell population dynamics in mammalian aging. Science 387, eadn3949 (2025). 10.1126/science.adn3949

31 Emont, M. P. et al. A single-cell atlas of human and mouse white adipose tissue. Nature 603, 926–933 (2022). 10.1038/s41586-022-04518-2

32 Sarvari, A. K. et al. Plasticity of Epididymal Adipose Tissue in Response to Diet-Induced Obesity at Single-Nucleus Resolution. Cell Metab 33, 437–453 e435 (2021). 10.1016/j.cmet.2020.12.004

33 Gonzalez-Hurtado, E. et al. Nerve-associated macrophages control adipose homeostasis across lifespan and restrain age-related inflammation. Nat Aging 5, 1828–1843 (2025). 10.1038/s43587-025-00952-9

34 Zechner, R. et al. FAT SIGNALS--lipases and lipolysis in lipid metabolism and signaling. Cell Metab 15, 279–291 (2012). 10.1016/j.cmet.2011.12.018

35 Wang, G. et al. Distinct adipose progenitor cells emerging with age drive active adipogenesis. Science 388, eadj0430 (2025). 10.1126/science.adj0430

36 Wagner, A. et al. Metabolic modeling of single Th17 cells reveals regulators of autoimmunity. Cell 184, 4168–4185 e4121 (2021). 10.1016/j.cell.2021.05.045

37 Ferrara, N. & Alitalo, K. Clinical applications of angiogenic growth factors and their inhibitors. Nat Med 5, 1359–1364 (1999). 10.1038/70928

38 Herold, J. & Kalucka, J. Angiogenesis in Adipose Tissue: The Interplay Between Adipose and Endothelial Cells. Front Physiol 11, 624903 (2020). 10.3389/fphys.2020.624903

39 Schoenborn, J. R. & Wilson, C. B. Regulation of interferon-gamma during innate and adaptive immune responses. Adv Immunol 96, 41–101 (2007). 10.1016/S0065-2776(07)96002-2

40 Saltiel, A. R. & Olefsky, J. M. Inflammatory mechanisms linking obesity and metabolic disease. J Clin Invest 127, 1–4 (2017). 10.1172/JCI92035

41 Xu, M. et al. JAK inhibition alleviates the cellular senescence-associated secretory phenotype and frailty in old age. Proc Natl Acad Sci U S A 112, E6301–6310 (2015). 10.1073/pnas.1515386112

42 Biran, A. et al. Quantitative identification of senescent cells in aging and disease. Aging Cell 16, 661–671 (2017). 10.1111/acel.12592

43 Acosta, J. C. et al. A complex secretory program orchestrated by the inflammasome controls paracrine senescence. Nat Cell Biol 15, 978–990 (2013). 10.1038/ncb2784

44 Huang, W., Hickson, L. J., Eirin, A., Kirkland, J. L. & Lerman, L. O. Cellular senescence: the good, the bad and the unknown. Nat Rev Nephrol 18, 611–627 (2022). 10.1038/s41581-022-00601-z

45 Birch, J. & Gil, J. Senescence and the SASP: many therapeutic avenues. Genes Dev 34, 1565–1576 (2020). 10.1101/gad.343129.120

46 Xu, M. et al. Senolytics improve physical function and increase lifespan in old age. Nat Med 24, 1246–1256 (2018). 10.1038/s41591-018-0092-9

47 Roche, P. A. & Furuta, K. The ins and outs of MHC class II-mediated antigen processing and presentation. Nat Rev Immunol 15, 203–216 (2015). 10.1038/nri3818

48 Amersfoort, J., Eelen, G. & Carmeliet, P. Immunomodulation by endothelial cells - partnering up with the immune system? Nat Rev Immunol 22, 576–588 (2022). 10.1038/s41577-022-00694-4

49 Pober, J. S., Merola, J., Liu, R. & Manes, T. D. Antigen Presentation by Vascular Cells. Front Immunol 8, 1907 (2017). 10.3389/fimmu.2017.01907

50 Jin, K. et al. Brain-wide cell-type-specific transcriptomic signatures of healthy ageing in mice. Nature 638, 182–196 (2025). 10.1038/s41586-024-08350-8

51 Collins, T. et al. Immune interferon activates multiple class II major histocompatibility complex genes and the associated invariant chain gene in human endothelial cells and dermal fibroblasts. Proc Natl Acad Sci U S A 81, 4917–4921 (1984). 10.1073/pnas.81.15.4917

52 Steimle, V., Siegrist, C. A., Mottet, A., Lisowska-Grospierre, B. & Mach, B. Regulation of MHC class II expression by interferon-gamma mediated by the transactivator gene CIITA. Science 265, 106–109 (1994). 10.1126/science.8016643

53 Islam, M. T. et al. Senolytic drugs, dasatinib and quercetin, attenuate adipose tissue inflammation, and ameliorate metabolic function in old age. Aging Cell 22, e13767 (2023). 10.1111/acel.13767

54 Minamino, T. et al. A crucial role for adipose tissue p53 in the regulation of insulin resistance. Nat Med 15, 1082–1087 (2009). 10.1038/nm.2014

55 Schafer, M. J. et al. Exercise Prevents Diet-Induced Cellular Senescence in Adipose Tissue. Diabetes 65, 1606–1615 (2016). 10.2337/db15-0291

56 Shiao, S. L., McNiff, J. M. & Pober, J. S. Memory T cells and their costimulators in human allograft injury. J Immunol 175, 4886–4896 (2005). 10.4049/jimmunol.175.8.4886

57 Shiao, S. L. et al. Human effector memory CD4+ T cells directly recognize allogeneic endothelial cells in vitro and in vivo. J Immunol 179, 4397–4404 (2007). 10.4049/jimmunol.179.7.4397

58 Carambia, A. et al. TGF-beta-dependent induction of CD4(+)CD25(+)Foxp3(+) Tregs by liver sinusoidal endothelial cells. J Hepatol 61, 594–599 (2014). 10.1016/j.jhep.2014.04.027

59 Taflin, C. et al. Human endothelial cells generate Th17 and regulatory T cells under inflammatory conditions. Proc Natl Acad Sci U S A 108, 2891–2896 (2011). 10.1073/pnas.1011811108

60 Lopes Pinheiro, M. A., et al. Internalization and presentation of myelin antigens by the brain endothelium guides antigen-specific T cell migration. Elife 5 (2016). 10.7554/eLife.13149

61 Oakley, R. & Tharakan, B. Vascular hyperpermeability and aging. *Aging Dis* **5**, 114–125 (2014). 10.14336/AD.2014.0500114

62 Xue, J. et al. Research Progress and Molecular Mechanisms of Endothelial Cells Inflammation in Vascular-Related Diseases. J Inflamm Res 16, 3593–3617 (2023). 10.2147/JIR.S418166

63 Reglero-Real, N., Rolas, L. & Nourshargh, S. Aging microvasculature: Effects on immune cell trafficking and inflammatory diseases. J Exp Med 222 (2025). 10.1084/jem.20242154

64 Long, J. Z., Lackan, C. S. & Hadjantonakis, A. K. Genetic and spectrally distinct in vivo imaging: embryonic stem cells and mice with widespread expression of a monomeric red fluorescent protein. BMC Biotechnol 5, 20 (2005). 10.1186/1472-6750-5-20

65 Barnden, M. J., Allison, J., Heath, W. R. & Carbone, F. R. Defective TCR expression in transgenic mice constructed using cDNA-based alpha- and beta-chain genes under the control of heterologous regulatory elements. Immunol Cell Biol 76, 34–40 (1998). 10.1046/j.1440-1711.1998.00709.x

66 Lewandoski, M., Meyers, E. N. & Martin, G. R. Analysis of Fgf8 gene function in vertebrate development. Cold Spring Harb Symp Quant Biol 62, 159–168 (1997).

67 Zhang, J. et al. An immune-based tool platform for in vivo cell clearance. Life Sci Alliance 6 (2023). 10.26508/lsa.202201869

68 Parlee, S. D., Lentz, S. I., Mori, H. & MacDougald, O. A. Quantifying size and number of adipocytes in adipose tissue. Methods Enzymol 537, 93–122 (2014). 10.1016/B978-0-12-411619-1.00006-9

69 Fleming, S. J. et al. Unsupervised removal of systematic background noise from droplet-based single-cell experiments using CellBender. Nat Methods 20, 1323–1335 (2023). 10.1038/s41592-023-01943-7

70 Wolf, F. A., Angerer, P. & Theis, F. J. SCANPY: large-scale single-cell gene expression data analysis. Genome Biol 19, 15 (2018). 10.1186/s13059-017-1382-0

71 Kalucka, J. et al. Single-Cell Transcriptome Atlas of Murine Endothelial Cells. Cell 180, 764–779 e720 (2020). 10.1016/j.cell.2020.01.015

72 Badia, I. M. P. et al. decoupleR: ensemble of computational methods to infer biological activities from omics data. Bioinform Adv 2, vbac016 (2022). 10.1093/bioadv/vbac016

73 Franzen, O., Gan, L. M. & Bjorkegren, J. L. M. PanglaoDB: a web server for exploration of mouse and human single-cell RNA sequencing data. Database (Oxford) 2019 (2019). 10.1093/database/baz046

74 Love, M. I., Huber, W. & Anders, S. Moderated estimation of fold change and dispersion for RNA-seq data with DESeq2. Genome Biol 15, 550 (2014). 10.1186/s13059-014-0550-8

75 Muzellec, B., Telenczuk, M., Cabeli, V. & Andreux, M. PyDESeq2: a python package for bulk RNA-seq differential expression analysis. Bioinformatics 39 (2023). 10.1093/bioinformatics/btad547

76 Chi, J., Crane, A., Wu, Z. & Cohen, P. Adipo-Clear: A Tissue Clearing Method for Three-Dimensional Imaging of Adipose Tissue. J Vis Exp (2018). 10.3791/58271

77 Dimitrov, D. et al. LIANA+ provides an all-in-one framework for cell-cell communication inference. Nat Cell Biol 26, 1613–1622 (2024). 10.1038/s41556-024-01469-w

78 Dobin, A. et al. STAR: ultrafast universal RNA-seq aligner. Bioinformatics 29, 15–21 (2013). 10.1093/bioinformatics/bts635

79 Putri, G. H., Anders, S., Pyl, P. T., Pimanda, J. E. & Zanini, F. Analysing high-throughput sequencing data in Python with HTSeq 2.0. Bioinformatics 38, 2943–2945 (2022). 10.1093/bioinformatics/btac166

80 Fang, Z., Liu, X. & Peltz, G. GSEApy: a comprehensive package for performing gene set enrichment analysis in Python. Bioinformatics 39 (2023). 10.1093/bioinformatics/btac757

81 Binder, J. X. et al. COMPARTMENTS: unification and visualization of protein subcellular localization evidence. Database (Oxford) 2014, bau012 (2014). 10.1093/database/bau012

82 Bellantuono, I. et al. A toolbox for the longitudinal assessment of healthspan in aging mice. Nat Protoc 15, 540–574 (2020). 10.1038/s41596-019-0256-1

83 Selahi, A. et al. Lymphangion-chip: a microphysiological system which supports co-culture and bidirectional signaling of lymphatic endothelial and muscle cells. Lab Chip 22, 121–135 (2021). 10.1039/d1lc00720c

